# Opposing Functions of Distinct Regulatory T Cell Subsets in Colorectal Cancer

**DOI:** 10.1101/2025.02.07.637083

**Authors:** Xiao Huang, Dan Feng, Sneha Mitra, Emma S. Andretta, Nima B. Hooshdaran, Aazam P. Ghelani, Eric Y. Wang, Joe N. Frost, Victoria Lawless, Aparna Vancheswaran, Qingwen Jiang, Christina S. Leslie, Alexander Y. Rudensky

## Abstract

Regulatory T (Treg) cells contribute to solid organ cancer progression, except in colorectal cancer (CRC) despite being abundantly present. Here, we demonstrate that two distinct tumoral IL-10⁺ and IL-10⁻ Treg cell subsets exert opposing functions by counteracting and promoting CRC tumor growth, respectively. The tumor restraining activity of IL-10⁺ Treg cells was mediated by their suppression of effector CD4 T cell production of IL-17, which directly stimulates CRC tumor cell proliferation. Consistently, IL-10⁻ Treg cells were more abundant in both mouse and human CRC tumors than in tumor-adjacent normal tissues, whereas IL-10^+^ Treg cells exhibited the opposite distribution. Furthermore, relative abundance of IL-10⁺ and IL-10⁻ Treg cells correlated with better and worse disease prognoses in human CRC, respectively. This functional dichotomy between Treg cell subsets provides a rationale for therapeutic strategies to selectively target pro-tumoral Treg cells while preserving their anti-tumoral counterparts across barrier tissue cancers that harbor both subsets.

## INTRODUCTION

Regulatory T (Treg) cells, expressing the transcription factor Foxp3, carry out a wide range of functions essential for organismal physiology and fitness in health and disease. By modulating innate and adaptive immune responses, they prevent fatal autoimmunity and inflammation while preserving tissue function by limiting both microbe-induced inflammation associated with host-commensal interactions or infections, and sterile metabolic inflammation. Additionally, Treg cells directly promote tissue repair during injuries by producing factors that support tissue maintenance and differentiation and by partaking in tissue stem cell niches.^1^ Parallel to the effector arm of the adaptive immune system, Treg cells patrol the organism through their recirculation and residence in the secondary lymphoid organs. In addition, they seed non-lymphoid tissues, particularly the barrier sites such as the gastrointestinal (GI) and genital tracts, skin, and airways. In these tissues, exposed to a variety of biotic and abiotic stressors, Treg cells establish and maintain immunological tolerance and limit inflammation through diverse immunosuppressive mechanisms.^2,3^

Both the tolerogenic and tissue-maintenance functions of Treg cells feature prominently in solid organ cancers, where they assume a highly activated state. Tumoral Treg cells are thought to regulate activities of other accessory cells of immune and non-immune origins in the tumor microenvironment (TME) or act on tumors directly to favor tumor progression. The pronounced enrichment of tumor-associated Treg cells in solid organ cancers, generally linked to a poor prognosis,^4–10^ has inspired major recent efforts towards development of strategies for targeting these cells as a novel means of immunotherapy. In support of these efforts, depletion of Treg cells in experimental solid organ cancer models, including PD-1 and CTLA-4 blockade-resistant genetically induced tumors and orthotopically transplanted cell lines derived from these tumors, resulted in their pronounced growth restraint or rejection.^11–13^ In contrast to the broad consensus on tumor-supportive functions of Treg cells in the vast majority of the established tumors, colorectal cancer (CRC), which ranks as the second leading cause of cancer-related mortality worldwide,^14^ appears to be an exception. In this regard, a recent study reported a positive correlation between Treg alongside CD8 T cell density in human CRC tumors and enhanced tumor control.^15^ CRC is generally classified into two primary types based on the microsatellite stability status and the proficiency of their mismatch repair (MMR): 80–85% of CRC cases are characterized by stable microsatellite DNAs (microsatellite stable; MSS) and proficient mismatch repair mechanisms (mismatch repair proficient; MMRp); the remaining 15% exhibit microsatellite instability (microsatellite instability-high; MSI-H) and deficient mismatch repair mechanisms (mismatch repair deficient; MMRd). Although blockade of PD-1 alone and in combination with CTLA-4 have recently shown impressive therapeutic responses in MSI-H CRCs, MSS CRCs remain by and large resistant to these interventions.^16,17^ A notable feature distinguishing the MSS CRC tumor microenvironment (TME) from the MSI-H CRC is the prevalence of Treg cells in the former.^15^ Mouse studies showed a high degree of heterogeneity of colonic Treg cells that stems from their ontogeny,^18–20^ differentiation states, and specific effector features.^21^ These effector features can contribute to different aspects of colon tissue biology as exemplified by a recent observation of terminally differentiated interleukin-10 expressing (IL-10^+^) Treg cells preventing colon shortening.^21^ These findings raise a possibility that while in MSS CRC some tumoral Treg cells have an expected tumor-promoting function, a distinct subset of tumoral Treg cells may exert a seemingly incongruous anti-tumoral function by restraining activity of tumor-supportive accessory cells.

Here, we tested this supposition by investigating the heterogeneity of transcriptional and chromatin states of tumoral vs. colonic Treg cells at a single-cell resolution in an orthotopic mouse model of MSS CRC that faithfully recapitulates human disease, as well as in MSS CRC samples from human patients. Analysis of the dynamics of tumoral Treg population during progression of tumors, established upon transplantation of *Apc^null^Trp53^null^ Kras^G12D^* cancer organoids,^22^ showed an enrichment in IL-10 non-expressing (IL-10^−^) Treg cells over IL-10^+^ Treg cells, whereas the exact opposite pattern was observed in the normal colon. Accordingly, targeted depletion of IL-10^+^ Treg cells increased the amounts of IL-17 produced by effector CD4 T cells, which by acting directly on CRC tumors promoted their growth, while selective loss of their IL-10^−^ counterparts resulted in a profound tumor shrinkage. Furthermore, the observed dichotomy between corresponding Treg cell subsets was conserved in human MSS CRCs, where a higher prevalence of IL-10^+^ and IL-10^−^ Treg cells correlated with improved and worsened survival outcomes, respectively. Notably, meta-analysis of published single-cell multiome datasets identified Treg subsets with similar features in human lung and skin cancers. Together, our studies suggest that in MSS CRC, and likely other barrier tissue malignancies, tumoral IL-10^+^ Treg cells may exert an indirect antitumoral function within TME by suppressing production of tumor-supportive mediators (such as IL-17) by other accessory cells, including effector T cells. These findings offer a rationale for developing therapeutic strategies to selectively target distinct tumor-promoting Treg cell subsets, while sparing tumor-opposing subsets, for immunotherapy of MSS CRC and other epithelial barrier cancers.

## RESULTS

### The AKP tumor is a faithful experimental model of human MSS CRC

To investigate the heterogeneity and function of Treg cells in MSS CRC, we sought to employ an orthotopic mouse CRC model that recapitulates properties of the human disease, including CRC driver mutations, anatomic features of tumor growth, TME immune cell composition, prominent Treg cell presence and resistance to PD-1 blockade based therapy. Thus, we adopted orthotopic transplantation of mouse CRC stem cell organoids derived from intestinal stem cells with the induced hallmark mutations of human CRC: *Apc* and *Tp53* gene loss and oncogenic Kras^G12D^ expression (AKP).^22^ Tumors morphologically resembling human CRC protruded into the lumen and were macroscopically detectable at 2–4 weeks while spontaneous metastases were observed in the draining mesenteric lymph nodes (mLNs) and the liver at 10 weeks after implantation of organoid cells into the cecal wall (Figure 1A). Notably, the AKP TME showed overall significantly reduced immune cell presence with an enrichment in Treg cells and macrophages, and diminished effector CD8 T cell activities in comparison to hypermutated MC38 colon adenocarcinoma, a commonly used mouse model of MSI-H CRC (Figures 1B–D).^23^ The observed features distinguishing these orthotopic mouse models closely aligned with those of corresponding human CRC types (Figure S1),^24^ highlighting the pronounced immunosuppressive character of the AKP tumor model.^6^ Contrary to antibody-mediated PD-1 blockage sensitivity of MC38,^25^ AKP tumors were resistant to this therapeutic modality, which resulted in a mild increase in Treg cells with no significant changes in tumor volume or CD8 T cell activation status (Figures 1E–H). This once again mirrored the therapeutic response to PD-1 blockade and the lack thereof in the corresponding human CRC types. Together, these initial experiments established the AKP tumor as a faithful model of the human MSS CRC.

**Figure 1.**
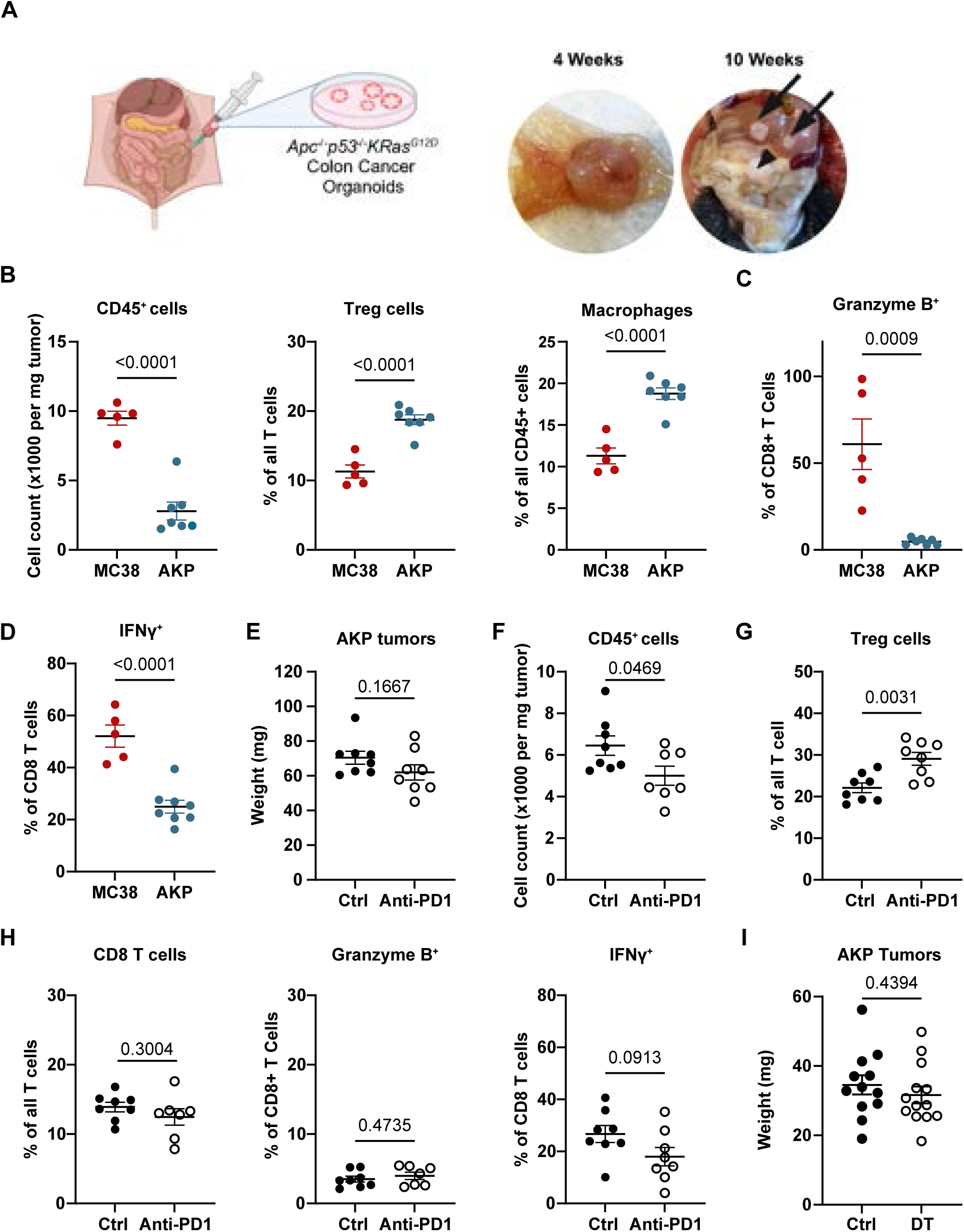
Orthotopic AKP tumor organoid model of human MSS CRC. (**A**) Left: the schematic of orthotopic AKP tumor organoid transplantation model; Right: representative images of primary and metastasized tumor at indicated time point. (**B–D**) The frequencies of indicated populations as measured by flow cytometric analysis of the indicated tumors 2 weeks after transplantation; (**E–F**) PD1 antibody was administered one week after tumor implantation for a period of two weeks. (**E**) tumor sizes at 2 weeks after indicated treatment; (**F**) the frequencies or absolute numbers of indicated populations in AKP tumors 2 weeks after indicated treatment. Data shown are pooled from 2–3 independent experiments.

### AKP tumor associated Treg and effector T cell heterogeneity

Considering that tumors regress upon Treg depletion in most experimental mouse models and that human MSS CRC seems deviant from a typical association of tumoral Treg “richness” with a poor prognosis, we investigated the efficacy of therapeutic Treg targeting in AKP tumor-bearing *Foxp3^DTR^* mice. In this mouse model, diphtheria toxin receptor (DTR) is expressed under the control of the endogenous *Foxp3* promoter, which enables punctual removal of Treg cells upon diphtheria toxin (DT) administration. In keeping with the reported lack of a clear link between Treg cell abundance and clinical disease progression in human CRC patients, we found that the DT-induced pan-Treg depletion did not significantly alter tumor growth (Figure 1I). While these results appeared to suggest that Treg cell functionality is dispensable for AKP tumor progression, it remained possible that distinct subsets within tumoral Treg cell population may exert opposing effects on tumor growth resulting in a zero-sum effect.

To address this possibility and to further substantiate the validity of the AKP tumor model for studying human disease, we first performed paired single-cell RNA/ATAC-seq analyses of highly purified T cell populations from primary AKP tumors and adjacent tumor-free cecal tissues 4 weeks after transplantation. After processing, quality control filtering, and clustering, 17 distinct clusters (15,621 cells) were identified based on the latent space representation of scRNA-seq data and visualized using uniform manifold approximation and projection (UMAP) (Figures 2A–2C). Analysis of the effector T cells showed that CD8 T cells encompassed three clusters of CD8αβ T cells, containing terminally differentiated (TD CD8, *Pdcd1*^hi^ *Gzmb*^lo^), effector (*Gzma*^hi^ *Gzmb*^hi^), and memory (*Il7r*^hi^ *Ccr7*^hi^) CD8αβ T cells, and one cluster of CD8αα T cells (Figure 2F). Effector CD4 T cells encompassed two Th1 (*Tbx21*^+^ *Ifng*^+^) clusters, alongside Th2 (*Gata3*^+^ *Il4*^+^ *Il5*^+^), Th17 (*Rorc*^+^ *Il17a*^+^ *Il17f*^+^), Tfh (*Cxcr5*^+^ *Bcl6*^+^), memory (*Ccr7*^hi^ *Sell*^hi^ *Il7r*^hi^), and recent draining LN emigrant (*S1pr1*^hi^, CD4_RLE_) clusters (one each). Accordingly, analysis of motif accessibility in ATAC-seq peaks in each cluster showed enrichment of corresponding lineage-defining transcription factor motifs (TBX for Th1 and CD8; TCF7 for memory cells; Gata for Th2s; Rorc for Th17) (Figure 2D). Beyond classical αβ T cells, we also identified two clusters of *Rorc*^+^ or cytotoxic *Gzma*^hi^ *Gzmb*^hi^ γδ T cell, as well as two recently described innate-like FcεR1γ^+^ T cells (ILTCKs) clusters.^26^ Most of the identified clusters contain both tumoral and colonic T cells, however, one cluster of highly activated PD-1^+^ Th1 cells appears to reside only in tumors (Figures 2C and 2F). Importantly, the effector T cell clusters identified in AKP tumors closely paralleled those observed in human MSS CRCs,^24^ lending further support for the exactitude of this model of human disease.

**Figure 2.**
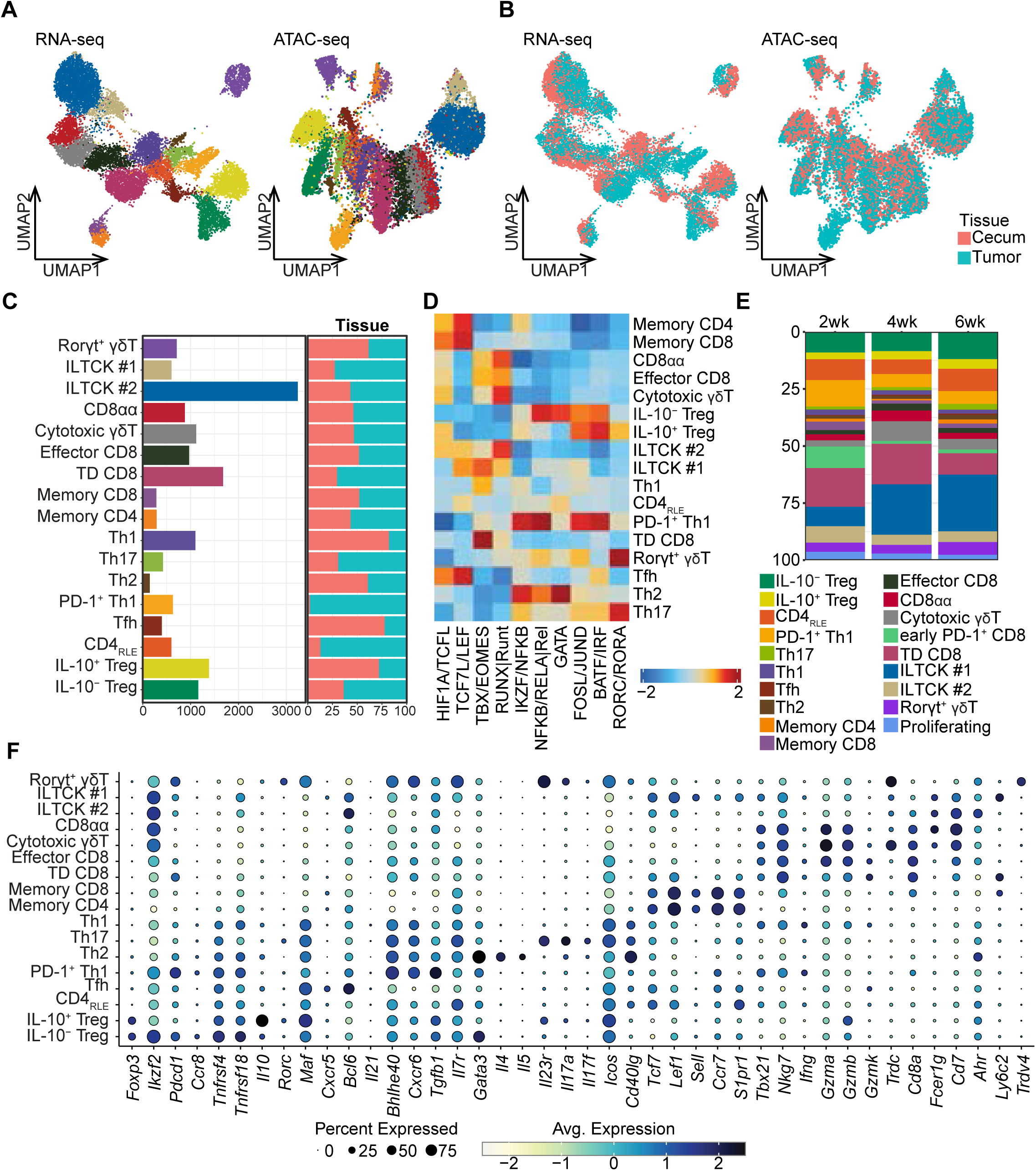
Paired single-cell RNA-seq and ATAC-seq analysis of CRC tumor associated T cells. (**A–B**) Paired scRNA-seq and scATAC-seq analysis of T cells isolated from AKP tumor or adjacent cecal tissues 4 weeks after tumor inoculation. UMAP plots are shown for (**A**) cell type annotations and (**B**) tissue annotations (n=15,621 cells). (**C**) Bar graphs depicting the number of cells for each cell type and the proportions for each tissue. (**D**) chromVAR motif enrichment analysis of multiome scATAC-seq data across different subsets of T cells. (**E**) The tumoral T cell subset dynamics 2, 4, and 6 weeks after tumor implantation determined by scRNA-seq (n=21,269 cells total: 10,013 from normal adjacent cecal tissues and 11,256 from AKP tumors). (**F**) The expression of selected genes of interest in AKP tumoral T cell subsets 4 weeks after tumor inoculation. TD CD8: terminally differentiated CD8 T cells. RLE CD4: recent LN emigrant CD4 T cells.

Treg cells formed two clusters, which were annotated as IL-10^+^ and IL-10^−^ Treg cells based on their expression of the *Il10* gene. While these two clusters were populated by both colonic and tumoral cells, the majority of IL-10^+^ Treg cells were found in the cecum. In contrast, the IL-10^−^ Treg cells were enriched in the tumor, raising a possibility of potentially distinct functions of IL-10^+^ and IL-10^−^ Treg cells in MSS CRC (Figure 2C). To gain insights into their dynamics during AKP tumor progression, we performed scRNA-seq of T cell populations from tumors and adjacent cecal tissues at 2, 4, and 6 weeks after AKP organoid implantation. All 17 clusters identified in the multiome analysis were captured by the time course scRNA-seq analysis, which also identified two additional early *Pdcd1*^hi^ CD8 and proliferative T cell clusters (Figure 2E). Extending the observation of the tumoral Treg subset composition bias revealed by single-cell multiome analysis, the scRNA-seq time course, corroborated by flow cytometric analysis, showed a notable enrichment in IL-10^−^ to IL-10^+^ Treg cells with tumor progression while the relative size of IL-10^+^ Treg cells did not significantly change (Figures 2E and S2B). The increase in IL-10^−^ Treg abundance was accompanied by a pronounced decline in tumor-specific PD-1^+^ Th1 cells and early *Pdcd1*^hi^ CD8 T cells. These two populations of effector cells were most abundant in early tumors at 2 weeks after implantation but not present in colonic tissues (Figures 2E and S2A). The observed increase in IL-10^−^ Treg cells aligns with their hypothesized tumor-promoting role, prompting further inquiry into the function of IL-10^+^ Treg cells.

### Divergent gene regulatory programs of tumor associated Treg cell subsets

The dynamics of IL-10^+^ and IL-10^−^ Treg cell subsets in CRC implied their distinct gene regulatory and transcriptional features. In this context, the former subset predominantly expressed *Rorc* gene, which encodes the transcription factor RORγt, whereas the latter expressed *Ikzf2* transcripts encoding Helios (Figure 2F). Thus, RORγt^+^ and Helios^+^ Treg cells can serve as surrogates for the corresponding IL-10^+^ and IL-10^−^ Treg cell subsets in the AKP tumors and adjacent normal colonic tissues (Figure 3A). To further dissect their divergent transcriptional regulation, we employed the SCARlink algorithm to identify putative enhancers of signature genes for both populations by analyzing paired ATAC- and RNA-seq datasets.^27^ Pseudo-bulk ATAC-seq tracks demonstrated differential accessibility between IL-10^+^ versus IL-10^−^ Treg cells at the putative enhancer sites in the *Il10* and *Ikzf2* loci, but not the *Foxp3* locus reflecting their distinct transcriptional regulatory circuitry (Figure 3B). Consistently, the difference in the number of gene-linked enhancers for some of the top differentially expressed genes closely correlated with the expression of the corresponding transcripts in these two subsets (Figure 3C).

**Figure 3.**
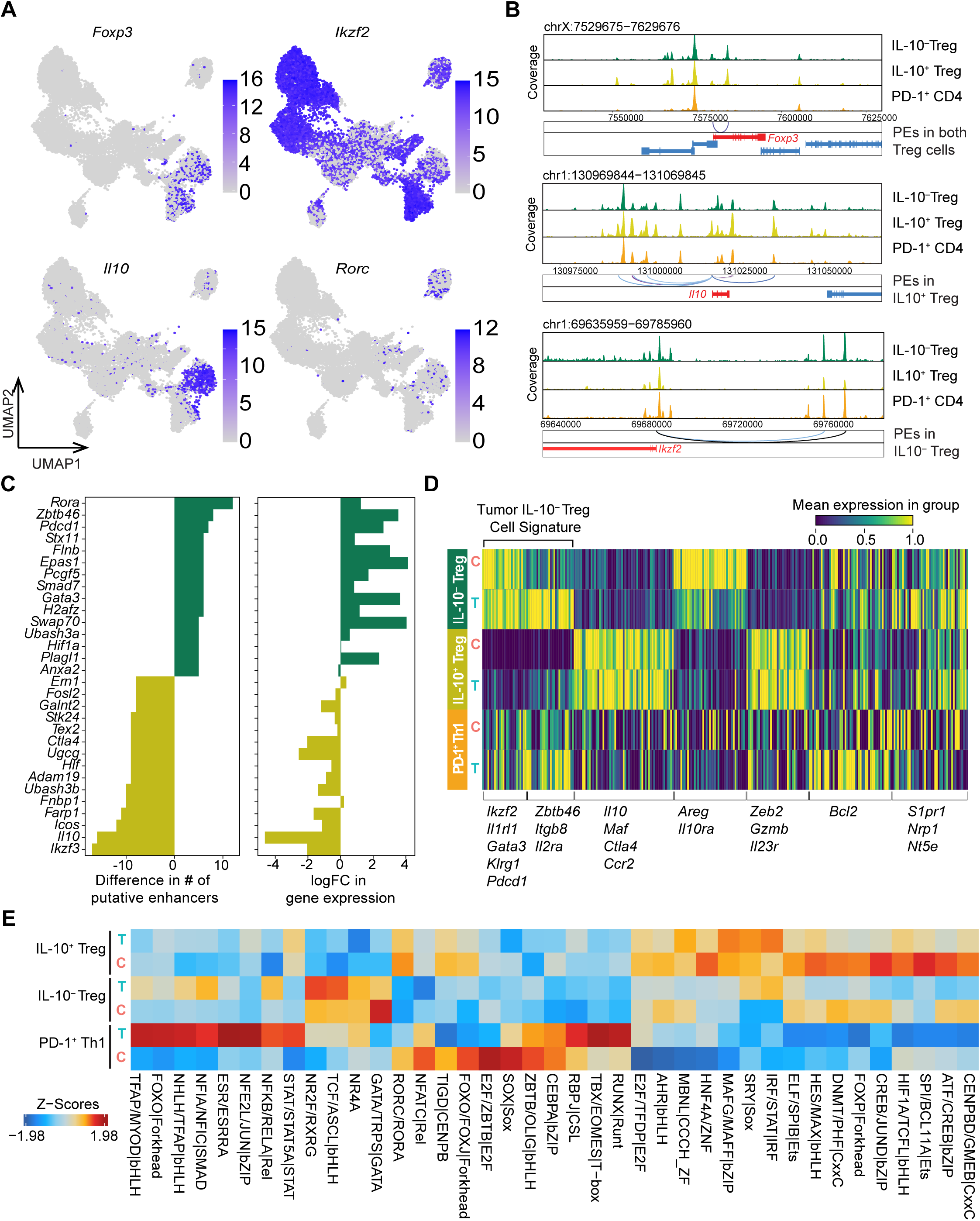
Heterogeneity of CRC tumoral Treg cells. (**A**) UMAP plots depicting the expression of marker genes distinguishing IL-10^+^ and IL-10^−^ Treg cells. (**B**) Gene browser chromatin accessibility tracks for *Foxp3*, *Il10*, and *Ikzf2* gene loci for IL-10^+^ and IL-10^−^ Treg cells. (**C**) Numbers of putative enhancers (left) and log fold change in gene expression (right) for select genes for IL-10^+^ and IL-10^−^ Treg cells. (**D**) Heatmap depicting expression of gene modules identified across all Treg subsets split by cells from tumor (T) and adjacent tumor-free cecum (C). (**E**) chromVAR analysis of differential motif enrichment for all Treg cell subsets with the indicated tissue origins (T, tumor; C, cecum).

To identify transcriptional modules expressed by IL-10^+^ versus IL-10^−^ Treg cells, we employed Hotspot, a graph-based method of neighborhood analysis.^28^ By calculating gene scores of the estimated modules, we identified those unique to each annotated Treg subset. Some of these modules also displayed tumor- or cecum-specific patterns indicating potential environmental imprinting of these regulatory programs (Figure 3D). Among the gene modules specific to IL-10⁻ but not IL-10⁺ Treg cells, two were enriched or shared in cells residing in AKP tumors. Henceforth referred to as the tumor IL-10⁻ Treg signature, these modules comprised either *Ikzf2*, *Gata3*, and *Klrg1* transcripts, or *Zbtb46*, *Itgb8*, and *Il2ra* transcripts. In contrast, another IL-10^−^ Treg specific module with characteristic *Areg* and *Il10ra* expression, was preferentially detected in cells residing in the adjacent normal colonic tissues. The observed expression pattern likely reflected the heightened states of activation of tumoral IL-10^−^ Treg cells and the tissue-supporting function of their colonic counterparts. While two separable gene modules were also identified as IL-10^+^ Treg cell-specific, their expression did not vary between the tumors and adjacent normal colonic tissues, indicating the lack of noticeable modulation of IL-10^+^ Treg cell transcriptional features by the CRC TME. These two IL-10^+^ Treg cell-specific modules were characterized by enrichment in *Il10, Maf, Ctla4,* and *Ccr2* transcripts, and *Zeb2, Gzmb,* and *Il23r* transcripts, respectively (Figure 3D). Analysis of the ATAC-seq data using chromVar showed that while IL-10^+^ and IL-10^−^ Treg cells shared certain TF binding motif enrichment at their unique accessible chromatin sites, the NR4A family motif was selectively enriched in tumoral IL-10^−^ Treg cells (Figure 3E). This result suggests a stronger TCR signaling experienced by the tumor-residing IL-10^−^ Treg cells in agreement with their heightened activation state implied by RNA-seq analysis (Figure 3D). In support of this notion, gene score analysis of tumoral IL-10^+^ Treg, IL-10^−^ Treg, and PD1^+^ Th1 cells showed the highest TCR signaling-related gene expression in tumoral IL-10^−^ Treg cells among all three subsets. (Figure S3). Collectively, these results are suggestive of functional divergence between IL-10^+^ and IL-10^−^ Treg in the AKP tumor model.

### IL-10^+^ and IL-10^−^ Treg cells have opposing functions in MSS CRC tumors

To directly investigate the functionality of IL-10^+^ and IL-10^−^ Treg cell subsets in the AKP tumor model, we employed their punctual selective depletion using two complementary genetic tools we developed. IL-10^+^ Treg cells were ablated upon DT administration into AKP tumor-bearing *Il10^tdTomato-Cre^ Foxp3^lsl-DTR^* (*Il10^Cre^Foxp3^lsl-DTR^*) mice, in which *Il10* locus encoded Cre recombinase excised a *loxP* site-flanked STOP cassette preceding the DTR coding sequence inserted into the *Foxp3* locus. Consequently, DTR is expressed exclusively in IL-10^+^ Treg cells in these mice. As a complementary approach, we induced selective DT-mediated ablation of IL-10^−^ Treg cells in AKP tumor-bearing *Il10^tdTomato-Cre^Foxp3^flox-DTR^*(*Il10^Cre^Foxp3^flox-DTR^*) mice, in which *Il10* locus encoded Cre excised the *loxP* site-flanked DTR coding sequence from the *Foxp3* locus, resulting in DTR expression exclusively in IL-10^−^ Treg cells. The experimental and control groups of mice were treated with DT or heat-inactivated DT (boiled DT; bDT), respectively, on day 1, 2, 5, 9, and 13 after intra-cecal implantation of AKP tumor organoids (Figure 4A). This resulted in efficient removal of DTR-expressing Treg cell populations in *Il10^Cre^Foxp3^lsl-DTR^* and *Il10^Cre^Foxp3^flox-DTR^* mice (Figure S4A), as indicated by the pronounced preferential depletion of tumoral RORγt^+^ and Helios^+^ Treg cells, respectively (Figures S4B–C). A moderate decrease in RORγt^+^ Treg cells observed in DT-treated *Il10^Cre^Foxp3^flox-DTR^* mice was likely due to the lack of *Il10* expression by some RORγt^+^ Treg cells. While these Treg cell subset perturbations did not significantly alter overall immune cell infiltration of AKP tumors (Figure S4D), IL-10^−^ Treg cell depletion led to a marked reduction in the tumor sizes, which upon closer histological and flow cytometrical examination showed negligible remaining tumor cell presence accompanied by pronounced increase in mononuclear cells and tumoral CD8 T cell responses. This observation suggests efficient mobilization of anti-tumor effectors upon removal of IL-10^−^ Treg cells (Figures 4C and S4E, bottom panels). In contrast, depletion of IL-10^+^ Treg cells resulted in a notable increase in tumor sizes without detectable changes in effector CD8 or CD4 T cell frequencies, suggesting the enhanced tumor growth was not a consequence of a diminished overall anti-tumoral T cell response, but its altered “flavor” favoring tumor progression (Figure S4E, top panels).

**Figure 4.**
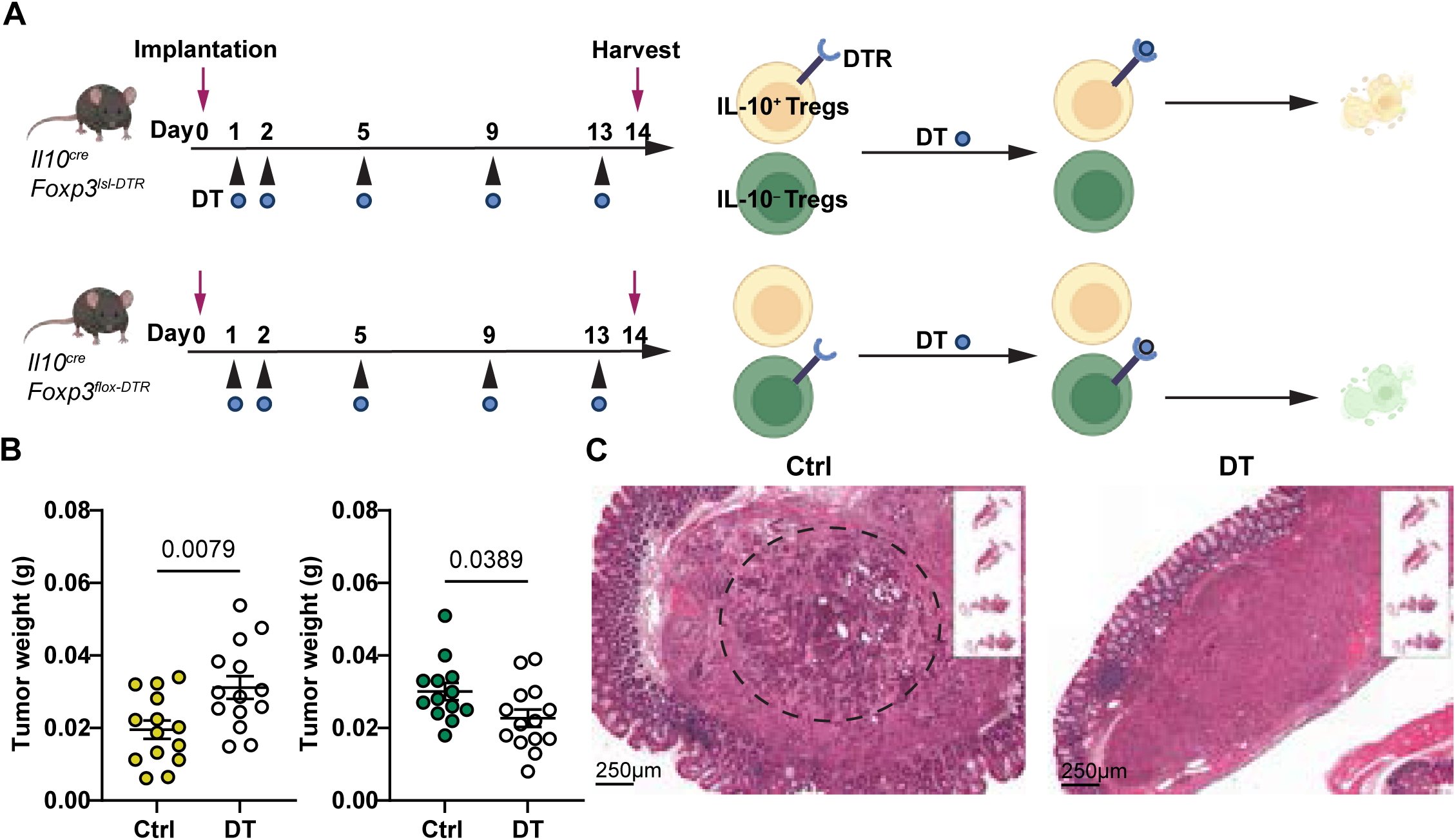
Targeted ablation of IL-10^+^ or IL-10^−^ Treg cells results in opposing effects on CRC tumor growth. (**A**) The experimental design schematic of diphtheria toxin (DT) induced selective ablation of IL-10^+^ and IL-10^−^ Treg cells using genetic mouse models. (**B)** Tumor sizes in DT- and control bDT-treated *Il10^tdTomato-Cre^ Foxp3^lsl-DTR^* (left) and *Il10^tdTomato-Cre^ Foxp3^flox-DTR^* mice (right) two weeks after AKP organoid implantation. Pooled data from two independent experiments are shown. (**C**) H&E staining of AKP tumors from indicated treatment groups. Images are representative of 3 independent experiments (dashed circle in the left panel indicates the tumor cell area).

To characterize potentially divergent effects of DT mediated depletion of these Treg cell subsets on the AKP TME, we first examined the myeloid populations in AKP tumors 2 weeks after implantation of tumor organoids in *Il10^Cre^Foxp3^lsl-DTR^* and *Il10^Cre^Foxp3^flox-DTR^* mice. Ablation of IL-10^+^ Treg cells resulted in a significant increase in Ly6C^hi^ monocytes, macrophages, and neutrophils, but not mast cells or eosinophils, indicative of a heightened type 3 response (Figures 5 and S5A, top panels). Tumor-associated macrophages and neutrophils are known for their tumor-promoting functions in CRC.^29–31^ In contrast, IL-10^−^ Treg cell ablation following tumor inoculation resulted in decreased monocyte, macrophage, and neutrophil abundances in the TME, accompanied by a notable increase in eosinophils, potentially reflective of an increased type 2 response (Figures 5 and S5A, bottom panels). Indeed, IL-10^+^ Treg depletion led to a pronounced increase in *Rorc* and type 3 cytokine (IL-17 and IL-22) expression by CD4 T cells with a concomitant decrease in type 2 cytokines (IL-4 and IL-13), whilst granzyme B or IFNγ expression by CD8 T cells was unchanged (Figures 6A–B, top panels; Figures 6C and S5B). In a conspicuous contrast, depletion of IL-10^−^ Treg cells led to an increase in IFNγ producing CD8 T cells and type 2 cytokine producing CD4 T cells coupled with a decrease in IL-17 expressing Th17 cells (Figures 6A–B, bottom panels). Together, these data demonstrated that in AKP tumors IL-10^+^ Treg cells selectively repress Th17 responses, whereas IL-10^−^ Treg cells repress Th2 and CD8 T cell responses. Collectively, these results suggest that IL-10^+^ and IL-10^−^ Treg cells regulate the CRC TME by modulating the tumoral immune activation “tone.”

**Figure 5.**
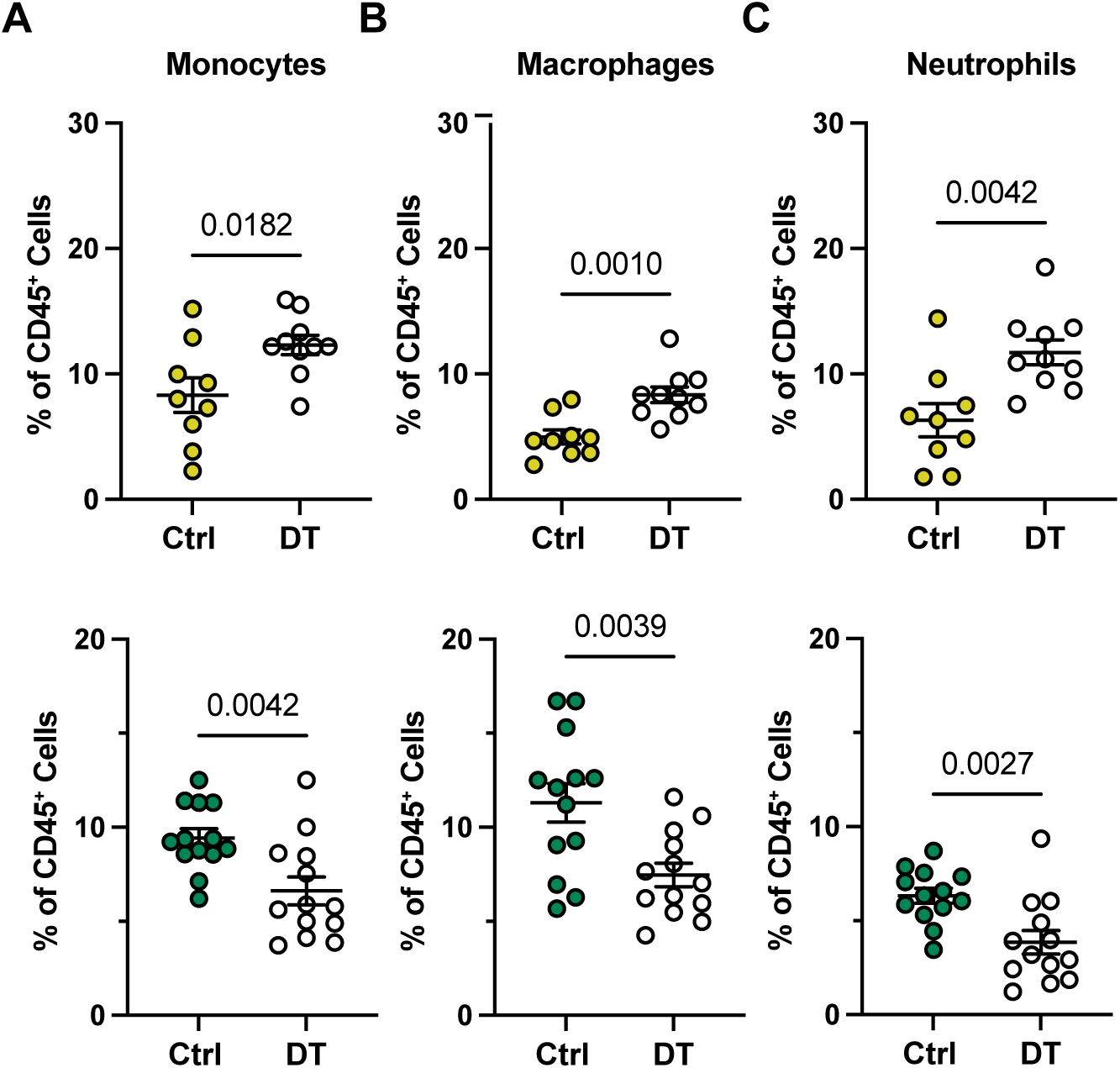
IL-10^+^ or IL-10^−^ Treg cell depletion causes contrasting shifts in CRC myeloid cell composition. (**A–C**) Flow cytometric analysis of the frequencies of monocytes (**A**), macrophages (**B**), and neutrophils (**C**) in AKP tumors from indicated treatment groups of *Il10^tdTomato-Cre^ Foxp3^lsl-DTR^* (top) and *Il10^tdTomato-Cre^ Foxp3^flox-DTR^* mice (bottom) two weeks after AKP organoid implantation. Pooled data from two independent experiments are shown.

**Figure 6.**
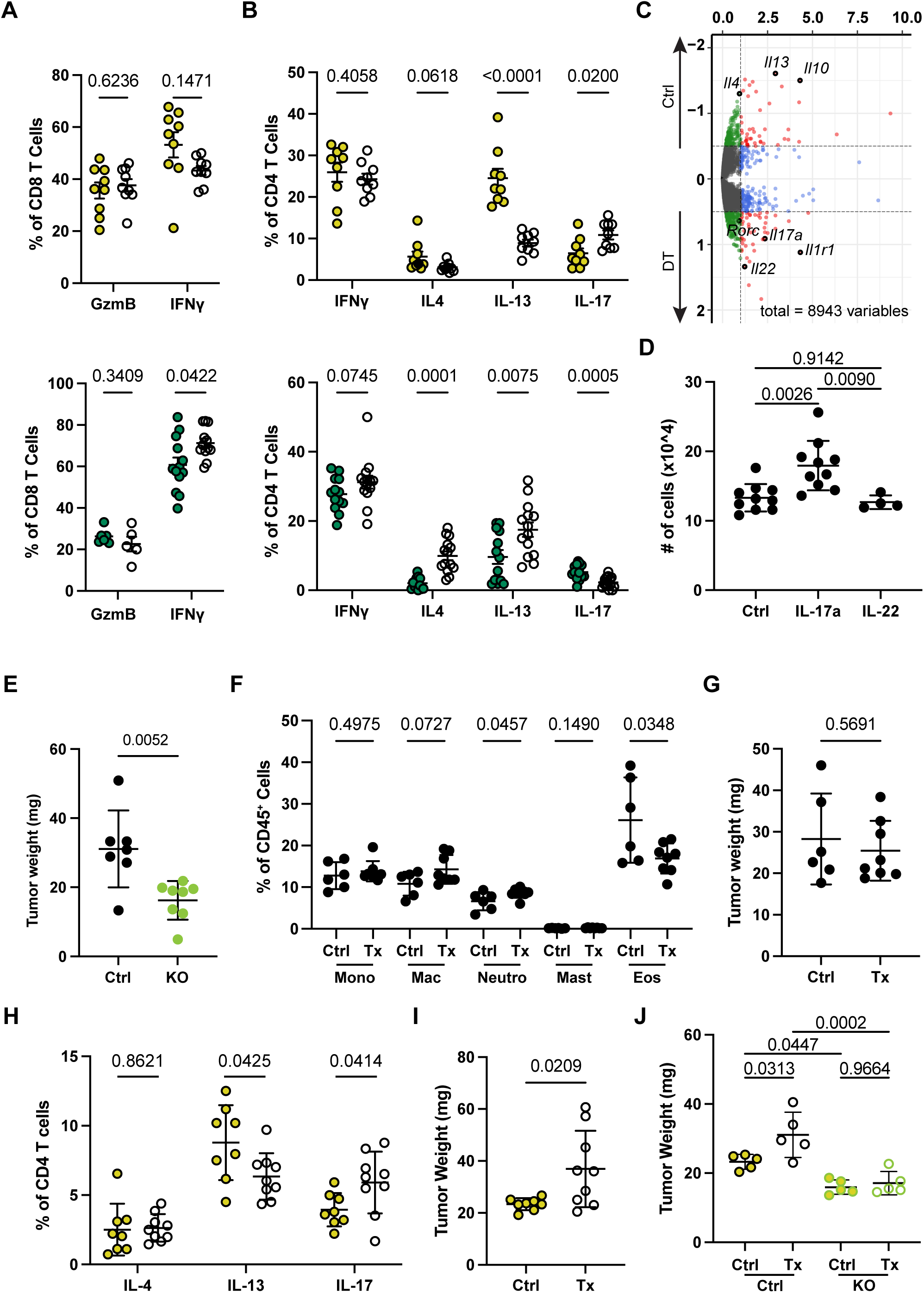
IL-10^+^ and IL-10^−^ Treg cells exert anti-and pro-tumoral functions through regulation of CD4 T cell responses. (**A–B**) Flow cytometric analysis of the indicated cytokines production by PMA/ionomycin stimulated CD8 (**A**) and CD4 T cells (**B**) isolated from AKP tumors from *Il10^tdTomato-Cre^ Foxp3^lsl-DTR^* (top) and *Il10^tdTomato-Cre^ Foxp3^flox-DTR^* mice (bottom) two weeks after their implantation. (**C**) Pseudo-bulk RNA-seq analysis of differential gene expression in tumoral CD4 T cells from DT vs control (bDT) treated *Il10^tdTomato-Cre^ Foxp3^lsl-DTR^* mice two weeks after AKP organoid implantation. Key genes of interest are labeled. (**D**) Frequencies and absolute numbers of CD8 and CD4 T cell numbers producing indicated effector molecules. (**E**) Weights of *Il17ra*-sufficient (WT) and -deficient (KO) AKP tumors two weeks after their implantation into C57BL/6 mice. (**F–I**) Flow cytometric analysis of the indicated cell types (**F, H**) and tumor weights (**G, I**) two weeks after the indicated treatments; for (**F)** and **(G)**, Tx: IL-4 and IL-13 neutralizing antibodies; Ctrl: isotype control IgG; for (**H)** and **(I)**, Tx: IL-10Rα blocking antibody; Ctrl: isotype control IgG. (**J**) Weights of *Il17r*-sufficient (WT) and -deficient (KO) AKP tumors two weeks after their implantation into C57BL/6 mice and indicated treatments (Ctrl, isotype control antibody injection; Tx, IL-10Rα blocking antibodies injection). Pooled data from two independent experiments are shown in all panels.

### IL-10^+^ Treg cells oppose CRC progression by limiting IL-17 production

Previous studies implicated IL-17 signaling in promoting early stages of transformation of intestinal epithelial cells in addition to vascularization and recruitment of pro-tumoral myeloid cells.^32–35^ Therefore, the increases in type 3 cytokines (IL-17 and IL-22) and tumor sizes noted upon depletion of IL-10^+^ Treg cells raised a question whether IL-17 or IL-22 could affect AKP tumors and whether the effect is direct or indirect. In support of a specific and direct activity of IL-17, we observed an increased growth of AKP tumor organoids *in vitro* in the presence of IL-17, but not IL-22 (Figure 6D). Analysis of the published bulk RNA-seq datasets revealed high *Il17ra* gene expression in similar mouse CRC organoids, lending further support for the idea that IL-17 may signal directly to AKP tumor cells to promote their proliferation.^36^ To test this hypothesis, we generated IL-17Rα*-*deficient AKP organoids using CRISPR-Cas9 mediated gene editing. Once transplanted into the cecum, the IL-17Rα-deficient tumors were of significantly smaller sizes than their control counterparts, implicating the direct signaling through IL-17Rα in IL-17-dependent growth of AKP tumors *in vivo* (Figure 6E). Because the enhanced tumor progression or its restraint caused by depletion of IL-10^+^ or IL-10^−^ Treg cells, respectively, were also associated with the opposing effects on IL-4 and IL-13 production, we tested whether a decrease in these cytokines could have contributed to increased tumor growth in the absence of IL-10^+^ Treg cells. While antibody mediated neutralization of IL-4 and IL-13 resulted in a marked reduction in tumoral eosinophils with a concomitant mild increase in tumoral neutrophils (Figure 6F), tumor sizes were unaffected in comparison to IgG isotype control treated group (Figure 6G). These data suggest that a decrease in type 2 cytokines is unlikely to contribute in a non-redundant manner to increased CRC tumor growth in the absence of IL-10^+^ Treg cells.

Previous studies showed that Th17 cells can be directly controlled through their IL-10R signaling in response to IL-10 produced by both Foxp3^+^ and Foxp3^−^ CD4 T cells.^37–39^ In light of these observations, our results suggest that IL-10^+^ Treg cell depletion accelerates tumor growth through IL-17R signaling in AKP cells, induced by increased IL-17 production once IL-10R-mediated restraint is lifted. To test this idea, we employed antibody-mediated blockade of IL-10Rα signaling in AKP tumor bearing mice. In corroboration of this supposition, IL-10Rα blockade resulted in an increase in IL-17 production by CD4 T cells and in tumor sizes similar to those observed upon IL-10^+^ Treg cell depletion (Figures 6H–I). Furthermore, IL-17Rα-deficient AKP tumors were insensitive to IL-10Rα blockade as their sizes were comparable to those in IgG isotype control-treated mice (Figure 6J). Taken together, our results suggest that while tumoral IL-10^−^ Treg cells expectedly promote CRC tumor progression, IL-10^+^ Treg cells restrain primary CRC tumors in an IL-10 signaling dependent manner. The latter limits production of IL-17, which acts directly on AKP organoids to promote their growth.

### The duality of Treg cell populations is conserved in human CRC

We next asked whether IL-10^+^ vs IL-10^−^ Treg cell subsets were present in human MSS CRC. To address this question, we performed paired multiomic scRNA/ATAC-seq analyses of total T cell populations isolated from surgically resected primary CRC tumors and matched normal adjacent colonic tissues (NATs) from 3 patients. Analysis of total 21,676 T cells, which formed 8 distinct clusters (Figures 7A–B and S6A; see Methods), showed Treg cells forming two clusters resembling mouse tumoral IL-10^+^ and IL-10^−^ Treg cells; one of these clusters preferentially expressed *IL10* and *RORC*, and the other *IKZF2* gene transcripts (Figures S6B–C). Similar to their differential abundance in normal tissue vs tumor in mice, the human IL-10^+^ Treg cells were enriched in NATs, while the IL-10^−^ Treg cells were enriched in tumor samples (Figure 7C). scATAC-seq data analysis using chromVar indicated similar TF motif enrichment patterns within chromatin accessible sites between corresponding mouse and human subsets, with the enrichment of RORC motif noted in IL-10^+^ Treg cells and GATA and NR4A motives in IL-10^−^ Treg cells (Figure 7D). To determine transcriptional similarity of human IL-10^+^ and IL-10^−^ Treg cells to their mouse counterparts, we employed mouse Treg cell subset-specific gene modules (Figure 3D) and determined their expression in human Treg cells. Both subsets exhibited significant enrichment for gene modules identified in their mouse counterparts (Figure 7E). Consistently, Hotspot analysis identified subset-specific gene modules in human CRC-infiltrating Treg cells that correspond to their mouse subsets. Notably, *IKZF2*, *GATA3*, *CCR8*, and *CADM1* were preferentially expressed by IL-10^−^ Treg cells, whereas *MAF* and *ZEB2* were preferentially expressed by IL-10^+^ Treg cells. Together, these results demonstrated that transcriptional and chromatin states of human CRC Treg cell subsets resembled those of mouse IL-10^+^ and IL-10^−^ Treg cells. Notably, our analysis of human CRC T cell datasets, made publicly available by 10X Genomics, also identified two tumor-infiltrating Treg cell subsets transcriptionally similar to the ones identified by our multiomic analysis (Figures S7A–E).^40^ The examination of the accompanying spatial transcriptomic (ST) datasets further corroborated the observations made in our tissue dissociation-based analysis by demonstrating an enrichment in IL-10^+^ Treg cells in NATs, while IL-10^−^ Treg cells were enriched in the tumor foci (Figures 6G and S7F and S7G). Of note, both Treg cell subsets seem to be preferentially localized to the tumor margins in the proximity of cancer-associated fibroblasts (Figures S7F and S7G).

**Figure 7.**
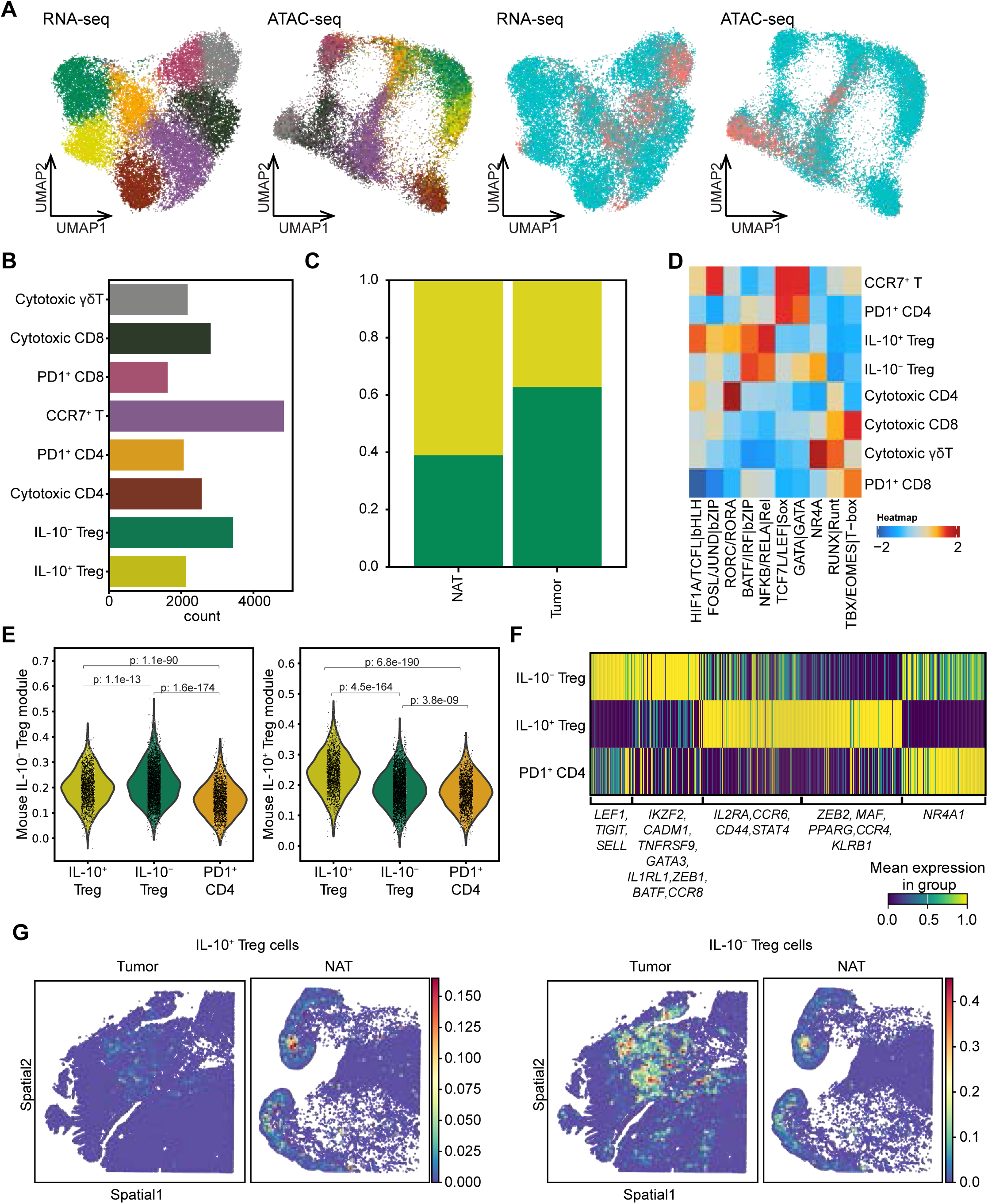
The IL-10^+^ and IL-10^−^ Treg cell subsets are present in human CRC. (**A**) Paired scRNA-seq and scATAC-seq analysis of T cells isolated from surgical samples of tumors and normal adjacent colon tissues (NATs) of 3 CRC patients. UMAP plots are shown for cell type annotations (left) and tissue annotations (right) for n=21,676 cells. (**B**) Bar graph depicting the number of cells for each cell type. (**C**) Proportions of IL-10^+^ and IL-10^−^ Treg cells in tumor and NAT. (**D**) chromVAR analysis of differential motif enrichment within scATAC-seq peaks across different subsets of T cells. (**E**) Gene scores calculated on the cells from human multiome analysis using the gene modules identified from mouse multiome analysis for IL-10^+^ and IL-10^−^ Treg cells in Figure 3D. (**F**) Heatmap depicting expression of gene modules identified across all Treg cells in tumors from CRC patients. (**G**) Probabilistic prediction of IL-10^+^ Treg cells (left) and IL-10^−^ Treg cells (right) in 10x VisiumHD spatial data from CRC tumor and NAT.^40^

Considering the opposing functions of IL-10^+^ and IL-10^−^ Treg cell subsets in tumor progression in mice, we next sought to investigate the potential association of their relative abundance in human CRC with clinical outcomes. We leveraged bulk RNA-seq datasets of surgical CRC samples from 102 patients and deconvolved these data using published scRNA-seq datasets for total CRC immune cells from 10X Genomics to infer relative abundances of the two Treg cell subsets (see Methods).^40^ The patients were then segregated into a high abundance group (H) and a low abundance group (L) based on their relative expression of characteristic gene signatures for each tumoral Treg cell population. Consistent with the anti-tumoral role of IL-10^+^ Treg cells established by mouse experiments, patients in the IL-10^+^ Treg cell high abundance group showed significantly better survival when compared to those in the corresponding low abundance group. Conversely, patients in IL-10^−^ Treg cell high abundance group exhibited significantly worse survival than in low abundance group, in agreement with the pro-tumoral function of this subset observed in mice (Figure S7H).

Having demonstrated the pro- and anti-tumoral functions of IL-10^+^ and IL-10^−^ Treg cells in mouse CRC and their association with better and worse human CRC prognosis, respectively, we next sought to determine whether similar populations of Treg cells can be identified in other human cancer types. Published single-cell atlas of T cells across 16 cancer types showed 7 distinct Treg cell clusters (Figure S8A).^41^ Upon closer examination, we found that clusters C0, C1, and C4 exhibited low expression for both IL-10^+^ and IL-10^−^ Treg cell gene modules, indicating their relatively less differentiated or activated states. Clusters C2, C3, and C5 showed high-level expression of IL-10^−^ Treg cell gene modules, while cluster C6 expressed a higher level of IL-10^+^ Treg cell signature module (Figures S8B and S8C). Although these patterns were indicative of overall transcriptional similarity of cluster 2, 3 and 5 cells to IL-10^−^ Treg cells, C3 and C5 cells were enriched only in basal cell carcinoma (BCC) and head and neck squamous cell carcinoma (HNSC), respectively, suggesting cancer type-dependent prevalence of a particular IL-10^−^ Treg cell flavor (Figure S8D). Importantly, both C2 and C6 populations showed significant presence in BCC, CRC, HNSC, and stomach adenocarcinoma suggesting that the dichotomy of tumoral IL-10^+^ and IL-10^−^ Treg cells is common across different barrier tissue cancers (Figure S8D).

## DISCUSSIONS

Initially characterized as the principal negative regulators of immune responses to “self” and “non-self” in the secondary lymphoid organs, Treg cells are increasingly recognized for their wide-ranging roles in non-lymphoid tissues. Treg cells contribute to tissue maintenance in two principal ways: indirectly, via suppression of tissue-damaging inflammatory responses; and directly, through production of factors that drive tissue differentiation, growth, and repair.^2,42^ Both direct and indirect functions encompass numerous molecular mechanisms such as production of immunomodulatory cytokine IL-10, consumption of IL-2, CTLA-4-assisted removal of CD80 and CD86 from dendritic cell surface, production of extracellular adenosine and epithelial growth factor family member amphiregulin (Areg), or activation of Notch signaling through expression of Notch ligand Jagged1.^2^ These mechanisms are mobilized through Treg cell recruitment, activation, proliferation, and differentiation in response to various stressors, including malignant transformation and tumor growth. While Treg cell activities can counteract precancerous hyperplasia and early tumor initiation by suppressing frank- or para-inflammation—potentially increasing the mutational load—they contribute to established tumor progression through indirect and direct mechanisms.^5^ Indirectly, Treg cells promote tumor growth by subduing cytotoxic and inflammatory responses of NK cells and tumor- or self-antigen-specific T cells, in conjunction with cell-intrinsic mechanisms, foremost PD-1 signaling, to drive these effector lymphocytes toward terminal differentiation and loss of pro-inflammatory function.^43,44^ The direct mechanisms may induce immediate signaling in tumor cells promoting their growth and fitness.^45^ Besides their association with progression of human cancers, increased numbers of activated tumoral Treg cells observed upon therapeutic PD-1 blockade have been linked to both cancer resistance and hyper-progression^46^ providing rationale for a combined PD-1 blockade and Treg depletion therapy.^11^

CRC is a notorious outlier in regard to the commonly accepted tumor-promoting function of Treg cells, with the conflicting reports on the correlation or lack thereof between Treg cells abundance and human CRC prognosis.^7–10,21^ One potential explanation for this discrepancy contends that the previous analyses did not discriminate between the contributions of Foxp3^hi^ Treg and Foxp3^lo^ effector T cells to overall *Foxp3* transcript amounts, and that the high numbers of the latter in CRC tumors correlate with microbial invasion and are linked to a better prognosis.^10^ However, effector T cell activation aside, microbiota and dietary antigens serve first and foremost as major drivers of heterogeneity of the colonic Treg cell population, stemming from extrathymic differentiation of RORγt^+^ and, to a lesser degree, Gata3^+^ Treg cells, in addition to activation of Treg cells of thymic origin residing in the colon.^18–20^

These findings motivated our investigation of intra-tumoral heterogeneity of Treg cells in CRC and its role in CRC progression. Our multimodal single-cell analyses demonstrated that IL-10^+^ RORγt^+^ and IL-10^−^ Helios^+^ Treg cells represent two transcriptionally and epigenetically distinct subsets of Treg cells residing in in the TME of mouse and human CRC and corresponding adjacent normal cecal tissues. These subsets exhibited differential enrichment of gene expression signatures downstream of TCR signaling,^47–49^ with the TCR-dependent gene signatures preferentially expressed in tumoral IL-10^−^ Treg cells but attenuated in IL-10^+^ Treg cells. These findings are consistent with the previous observations that withdrawal from TCR signaling is linked to the terminal differentiation of colonic IL-10^+^ Treg subset.^21,50–53^

In addition to their divergent transcriptomic and epigenomic features, opposing functions exerted by IL-10^+^ and IL-10^−^ Treg cells counteracting and promoting growth of CRC tumors, respectively, were revealed by selective ablation of individual subsets. This functional duality was consistent with the inversion in the relative abundance of these subsets in the tumors (% IL-10^+^ Treg < % IL-10^−^ Treg cells) in comparison to normal cecal tissues (% IL-10^+^ Treg > % IL-10^−^ Treg cells) observed in mice. Notably, similar trend was revealed by our analysis of human CRC and adjacent normal tissue ST datasets. This tumoral IL-10^+^ Treg dynamics could be due to progressively diminishing amounts of IL-2, resulting from its increased consumption by Treg cells and diminished production by effector T cells as CRC tumors grow.

The seemingly paradoxical function of IL-10^+^ Treg cells was predominantly due to their ability to suppress production of IL-17 by Th17 cells. Previous studies implicated IL-10 as a non-redundant control mechanism of Th17 cells, acting directly through IL-10R expressed by these cells.^37^ Fittingly, Treg cells express the highest amount of IL-10 on a per cell basis among IL-10 expressing cell types and account for the majority of IL-10 expressing T cells in the colon.^21^ In support of the above notion, antibody mediated blockade of IL-10R signaling resulted in a similar increase in IL-17 production and enhancement of the tumor growth. IL-17 is known to exert diverse effects on a wide range of cell types at barrier and other tissues. It mobilizes microbial defenses through the recruitment of neutrophils and monocytes and the induction of antimicrobial peptides. IL-17 also promotes fibrosis via augmenting production of extracellular matrix components, and induces adipose thermogenesis.^54^ Expectedly, a boost in IL-17 production observed upon relief of Th17 cells from IL-10^+^ Treg cell-mediated suppression was associated with increases in tumor-associated neutrophils, monocytes, and macrophages, the three cells type previously linked to CRC progression.^29–31^ While we have not formally excluded potential contribution of the latter to the observed enhanced AKP tumor growth, we found that contrary to IL-17Rα-sufficient tumors, which grew bigger upon IL-10Rα blockade, the IL-17Rα-deficient CRC size remained unaffected despite the increased IL-17 production. While Th17 cells also exhibited increased expression of IL-22, an epithelial maintenance factor promoting dextran sulfate sodium and azomethane induced colon tumorigenesis,^55^ we found IL-22 signaling in AKP tumors dispensable for boosting their growth, contrary to a non-redundant role of IL-17. Together, our results suggest that IL-10^+^ Treg cells restrain CRC progression by suppressing Th17 cells and thereby limiting amount of IL-17, which promotes AKP tumor cell growth.

The tumor-restraining activity of IL-10^+^ Treg cells was counterbalanced by tumor-promoting activity of IL-10^−^ Helios^+^ Treg cells, which was most likely linked to their superior ability to subdue tumoral CD8 T cell responses. While IL-10^−^ Treg cells also suppressed production of type 2 cytokines IL-4 and IL-13 by tumoral effector CD4 T cells and consequent recruitment of eosinophils to the TME, neutralization of these cytokines did not have any measurable effect on CRC growth. Notably, both mouse and human IL-10^−^ Treg cells but not IL-10^+^ Treg cells express high level of chemokine receptor CCR8. Previously, we and others have identified CCR8 as highly expressed by Treg cells across different human cancers.^4,56^ Subsequent studies revealed efficacy of CCR8 depleting antibody for Treg targeting as a monotherapy and in combinations with PD-1 or VEGF blockade or vaccination in experimental mouse and human cancer models.^57–59^ These results suggest that in CRC, CCR8 targeting-based therapeutic strategies may relieve suppression of effector antitumoral T cell responses while preserving suppression of production of tumor promoting factors by distinct Treg subsets. The relevance of the observed functional heterogeneity in our mouse studies to human MSS CRC is reflected in the association of an enrichment in gene expression signatures of human counterparts of mouse anti-tumoral IL-10^+^ and pro-tumoral IL-10^−^ Treg cells with a better and worse prognosis, respectively. Furthermore, our analysis of publicly available pan-cancer scRNA-seq datasets identified the counterparts of the two principal Treg cell subsets across other barrier tissue cancer types, suggesting potential broader clinical implications of the observed Treg cell heterogeneity beyond CRC.

In conclusion, our studies suggest that while IL-10^−^ Treg cells facilitate CRC tumor growth, IL-10^+^ Treg cells restrain tumor progression by quelling Th17 production of IL-17 that directly drives CRC tumor cell proliferation. On a more general note, the described Treg-mediated suppression of a production of tumor growth factor by another cellular component of TME represent a previously unappreciated cancer progression regulation mechanism. The functional dichotomy of the two major Treg cell subsets revealed by our studies of CRC offers rationale for the development of focused therapeutic strategies targeting pro-tumoral Treg cells while sparing their anti-tumoral counterparts in barrier tissue cancer types harboring both Treg subsets.

## Resource availability

### Lead contact

Further information and requests for resources and reagents may be directed to and will be fulfilled by the lead contact, Alexander Y. Rudensky.

### Materials availability

All reagents and resources used in this study should be directed to the lead contact. All reagents will be available based on requesting the lead contact and completing a Materials Transfer Agreement.

### Data and code availability

All sequencing datasets available on the GEO database with the following accession number: pending.

## ACKNOWLEDGMENTS

We thank the Memorial Sloan Kettering Integrated Genomics Core facility for performing all sequencing and the Single Cell Research Initiative for processing the scRNA-seq samples. We thank Dr. Scott Lowe for kindly providing us with the AKP tumor organoid. We thank Dr. Francisco Sanchez-Vega, Fan Wu, and Dr. Julio Garcia-Aguilar for providing us access to the bulk RNA-seq data from 102 patients. We thank Chin-Tung Chen and Dr. Julio Garcia-Aguilar for their assistance in obtaining patient tumor and normal adjacent tissue samples. This work was supported by the NCI Cancer Center Support Grant P30 CA008748, NIH grant R01 AI034206 (A.Y.R.), and the Ludwig Center at the Memorial Sloan Kettering Cancer Center (A.Y.R.). A.Y.R. is an investigator with the Howard Hughes Medical Institute. X.H. was supported by the Irvington Postdoctoral Fellowship from Cancer Research Institute and is supported by the Basic and Translational Immunology Postdoctoral Award from Ludwig Center for Cancer Immunotherapy. D.F. was supported by the institutional T32 grant at Memorial Sloan Kettering Cancer Center. The content is solely the responsibility of the authors and not necessarily represent the official views of the NIH.

## AUTHOR CONTRIBUTIONS

X.H., D.F., S.M., C.S.L., and A.Y.R. conceived the project. X.H. and D.F. designed, performed, and analyzed experiments with assistance from E.S.A., N.B.H., A.P.G., E.Y.W., J.N.F., V.L., A.V, and Q.J. S.M. and C.S.L. performed analysis of the single-cell sequencing experiments. X.H. and A.Y.R. wrote the manuscript with input from D.F., S.M., and C.S.L.

## METHODS

### Mice

*Foxp3^DTR^*, *Il10^tdTomato-Cre^ Foxp3^lsl-DTR^*, and *Foxp3^flox-DTR^*mice have been previously described.^21,60,61^ *Il10^tdTomato-Cre^ Foxp3^flox-DTR^*mice were generated by intercrossing. Co-housed littermates were used in all experiments, and all genotypes were represented in each litter analyzed. All mice were housed at the Research Animal Resource Center for Memorial Sloan Kettering Cancer Center (MSKCC) and Weill Cornell Medicine. All studies were performed under the protocol 08-10-023 and approved by the Sloan Kettering Institute Institutional Animal Care and Use Committee. All animals used in this study had no previous history of experimentation and were naive at the time of analysis.

### AKP tumor inoculation

AKP tumor organoid were maintained as previously described.^22^ For each experiment, tumor organoid 3D culture was digested with TrypLE and resuspend in PBS:Geltrex (2:1) mixture at the density of 10-20 × 10^6^ cells/ml and maintained on ice. 20ul of the cell suspension was injected into the subserosa space of mouse cecum.

### AKP tumor *in vitro* cytokine treatment

AKP tumor organoid were cultured *in vitro* as previously described.^22^ On day 1, cells were seeded at 0.67 million cells/ml in PBS:Geltrex (1:2) mixture and plated into 40 μl droplets at 1 drop per well concentration in a 24 well plate. Cells were treated with IL-17a and IL-22 at a concentration of 50 ng/ml each on day 1 and day 3. Cells from each well are harvested and counted on day 7.

### Gene editing of AKP tumor organoid

AKP organoids were electroporated using previously described protocol.^62^ 2μg of *in vitro* transcribed Cas9 mRNA, 200 pmol of sgRNAs (5’: GTGTTGATTACCGATGAGAA; 3’: GCAGCTAATCACCCCTAAGG) each, and 2μg of linearized donor plasmid were used per electroporation. Donor plasmids were designed with 40bp homology arms flanking both sgRNA cut sites. A targeting construct consisted of a EF1α promoter driving the Thy1.1 coding sequence, followed by a STOP codon, and polyA sequence was inserted into the *Il17ra* locus by microhomology-mediated end joining. The success of the knock-in was verified with genotyping PCR (Il17ra-F: GACTACTTGGCAGCAGAGCA; Il17ra-R: GCCAAGCCAGCACCACTG; EF1a-R: ACTTTCCCAGTTTACCCCGC) and flow cytometric analysis of Thy1.1 reporter expression. Organoid cells were FACS sorted based on their Thy1.1 expression twice before inoculation. For control organoid, only Cas9 RNA was used for the electroporation process.

### Mouse treatments

For diphtheria toxin (DT) treatment, DT was reconstituted in sterile PBS at 1 mg/ml and frozen at –80°C in single use aliquots. Aliquots were thawed and diluted 200 x in PBS. For inactivated control (bDT), this 1 ml dilution was heated at 95–100°C for 30 minutes. Both active and control DT were filtered through 0.22 μm syringe-driven filters. Mice were injected intravenously with 200 μl of this dilution for the first doses (1000 ng DT) and 200 μl of a 1:1 dilution with PBS for the second dose (500 ng DT) or intraperitoneally with 200 μl of a 1:1 dilution with PBS for the second dose for subsequent doses (500 ng DT).

For antibody treatment in tumor-bearing mice, the first dose immediately after the surgery was given by retroorbital i.v. injection, while all the subsequent injections were given i.p. PD1 antibody was injected on day 8, 11, 14, 17, and 20 at a dosage of 0.25 mg/mouse in 200 μl of saline. IL-10Rα antibody was injected on day 1 and day 8 after surgery at a dosage of 1 mg/mouse in 100 μl of saline. IL-4 and anti-IL-13 antibodies were injected on day 1, 4, 7, 10, and 13 at a dosage of 50 μg/mouse each in 200ul of saline.

### Cell isolation for flow cytometry

Mice were injected retro-orbitally with 1.5 μg anti-mouse CD45.2 (Brilliant Violet 510 conjugated, BioLegend 109838) in 200 μl sterile PBS 3 min prior to euthanasia to label and exclude blood-exposed cells. All centrifugations were performed at 700 x g for 3 min at 4 °C. Secondary lymphoid organs were dissected and placed in 1 ml wash medium (RPMI 1640, 2% Fetal Bovine Serum (FBS), 10 mM HEPES buffer, 1% penicillin/streptomycin, 2 mM L-glutamine). Tissues were then mechanically disrupted with the back end of a syringe plunger, and then passed through a 100 μm, 44% open area nylon mesh. For tumor and cecum, the cecal pouch was dissected, and after removal of fat and the cecal patch, opened longitudinally and vigorously shaken in 1x PBS to remove luminal contents. Tumor was then dissected from the surrounding cecal tissue. Both tumor and cecum were placed in a 50 ml screw-cap tube with 25 ml wash medium supplemented with 5 mM EDTA and 1 mM dithiothreitol and shaken horizontally at 250 RPM for 15 to 20 min at 37°C. After a 5 sec vortexing, epithelial and immune cells from the epithelial layer were removed by filtering the suspension through a tea strainer. Remaining tissue was placed back in 50 ml tubes, washed with 25 ml wash medium, strained again, and replaced in 5 ml tubes. Tumor was minced into pieces with the size of approximately 1–2 mm diameters. 25 ml wash medium supplemented with 0.2 U/ml collagenase A, 4.8 mM calcium chloride, 1 U/ml DNase I was added, along with four ¾ inch ceramic beads, and tissues were shaken horizontally at 250 RPM for 35 min at 37°C. Suspension was then passed through a 100 μm strainer, centrifuged to remove debris and collagenase solution, and then washed by centrifugation in 40% Percoll^TM^ in wash medium. All enzymatically digested samples were washed by centrifugation in 5 ml wash medium.

### Flow cytometry

To assess cytokine production after ex vivo restimulation, single cell suspensions were incubated for 4 hours at 37°C with 5% CO2 in the presence of 50 ng/ml PMA and 500 ng/ml ionomycin with 1 μg/ml brefeldin A and 2 μM monensin to inhibit ER and Golgi transport. For flow cytometric analysis, cells were stained in 96 well V-bottom plates. All centrifugations were performed at 700x g for 4 min at 4 °C for live cells and 900x g for 2 min for fixed cells. Staining with primary antibodies was carried out in 100 μl for 25 min at 4°C in staining buffer (PBS, 0.2 % (w/v) BSA, 2 mM EDTA, 10 mM HEPES, 0.1% (w/v) NaN3). Cells were then washed with 200 μl PBS and then concurrently stained with Zombie NIR™ Fixable Viability dye for 10 min at room temperature. Cells were washed with 100 μl staining buffer, resuspended in 200 μl staining buffer, and passed through a 100 μm nylon mesh. For cytokine staining, cells were fixed and permeabilized with BD Cytofix/Cytoperm per manufacturer instructions. Intracellular antigens were stained overnight hour at 4°C in the 1x Perm/Wash buffer. Samples were then washed twice in 200 μl 1x Perm/Wash buffer, resuspending each time, resuspended in 200 μl staining buffer, and passed through a 100 μm nylon mesh. All samples were acquired on an Aurora cytometer (Cytek Biosciences) and analyzed using FlowJo v10 (BD Biosciences).

### Cell sorting

For mouse samples, cell isolation was performed as described above, except that mice were perfused, and samples were not washed with 40% Percoll^TM^. Staining was performed as described above, except buffer contained 2 mM L-glutamine and did not contain NaN3, with staining volume adjusted to 500 μl and washes adjusted to 5 ml and staining performed in 15 ml screw-cap tubes. 1 μg ‘HashTag’ antibodies (BioLegend) were added to extracellular antigen stain for scRNA-seq sorting, and scRNA-seq sort samples were not DNase I treated. Samples were resuspended in Wash buffer supplemented 5mM EDTA for sorting and sorted into wash medium in 1.5 ml Protein LoBind tubes. For human CRC tissues, samples were weighed and dissociated with Tumor Dissociation Kit, Human (Miltenyi Biotec) on a gentleMACS^TM^ dissociator (Miltenyi Biotec) using programs recommended for CRC samples. Suspension was then passed through a 100 μm strainer, centrifuged to remove debris and collagenase solution, and then washed by centrifugation in RPMI extensively. All sorting was performed on an Aria II (BD Biosciences) or an AuroraCS cytometer (Cytek Biosciences).

### Histological analysis

The entire tumor tissue with ∼0.5 cm of surrounding cecal tissue were fixed in 4% PFA for > 48 hours. Tissue embedding, sectioning, and staining were carried out by the Molecular Cytology Core at MSKCC.

### Single-cell sequencing data preprocessing

The FASTQ files from all the single-cell datasets were aligned using the Cell Ranger pipeline. We performed quality control and applied different filtering thresholds for each dataset. For the mouse scRNA-seq data from the single-cell multiome of RNA+ATAC at 4 weeks, we restricted the percentage of mitochondrial reads (pct_counts_mt) to less than 10%, the number of genes expressed in each cell (n_genes_by_counts) to a maximum of 3,000, and the UMI counts (total_counts) to at least 1,000. For the scATAC-seq data from the same multiome, we retained cells with at least 180 fragments and a TSS enrichment score of at least 3. The TSS enrichment score is the ratio of the number of reads at the gene TSS relative to the flanking regions.^63^ For the mouse scRNA-seq data from 2, 4, and 6 weeks, we retained cells with pct_counts_mt less than 20%, n_genes_by_counts at most 3,000, and total_counts at least 500. In case of the metastasis scRNA-seq data from 10 weeks, we retained cells with pct_counts_mt less than 7%, n_genes_by_counts less than 4,200, and total_counts greater than 500. Finally, for the scRNA-seq from the multiome of patient samples, we extracted cells with pct_counts_mt less than 20%, n_genes_by_counts greater than 800, and total_counts less than 6,000. The matched scATAC-seq data from the patient samples were filtered to retain cells with at least 1,000 fragments and a TSS enrichment score of at least 4. For the multiome data sets, we retained the cells that passed quality control (QC) thresholds in both scRNA-seq and scATAC-seq.

### Batch correction and cell-type annotation

The single-cell multiome of the patient samples had strong batch effects in both scRNA-seq and scATAC-seq. To this end, we used combat^64,65^ on the log-normalized counts from scRNA-seq to generate batch-corrected counts. Using the batch-corrected counts, we performed principal component analysis (PCA) that we used to generate a UMAP embedding for visualization. We ran CellSpace^66^ on the top 25,000 variably accessible 500bp tiles with the margin parameter set to 0.1 to generate a batch corrected latent space embedding of the scATAC-seq data. This latent space embedding was used to generate a UMAP embedding for visualization. All the other remaining data sets did not have batch effects. In those cases, we performed dimensionality reduction using PCA on the normalized scRNA-seq counts and iterative LSI^63^ on the scATAC-seq tiles followed by UMAP generation. We performed Leiden clustering on the PCA components of scRNA-seq for all the data sets. The clustering was performed in an iterative manner where we sub-clustered specific clusters to obtain clearer signal of specific phenotypes. The cluster labels were transferred to scATAC-seq data for chromatin accessibility-based analyses.

### Transcription factor binding activity analysis

Peaks were called in the scATAC-seq data sets using Macs2^67^ by grouping the cells into pseudo-bulks based on the cell-type annotations obtained from matched scRNA-seq. chromVAR^68^ was run on these peak sets to quantify the enrichment of TF binding sites in the accessible regions of individual cells.

### Prediction of regulatory regions from single-cell multiome

SCARlink^27^ was run on the subsets of CD4 T cells from the mouse single-cell multiome. We trained models on 4,315 genes that were highly variable with detectable transcripts in at least 10% of all the CD4 T cells and extracted the predicted gene-linked enhancer tiles with z-score greater than 0.5 and false discovery rate (FDR) of 0.05 to compare the number of gene-specific enhancers predicted for IL-10^+^ and IL-10^−^ Tregs.

### Identification of gene modules

Hotspot^28^ was used separately on mouse and human scRNA-seq data to identify gene modules for Tregs. First, cells that were not Treg cells were filtered out alongside with genes that were not expressed in any of the Treg cells. Malat1/MALAT1, mitochondrial genes, and ribosomal genes were also filtered out. k-nearest neighbor (kNN) graphs were generated with 30 neighboring cells. For the mouse data, we generated modules with at least 25 genes at FDR < 0.05 and with at least 35 genes at FDR < 0.0001 for human data. This gave us seven gene modules in the mouse data and five gene modules in the human data. The gene scores calculated using these gene modules were generated using the score genes() function from scanpy.^65^

### Analysis of single-cell flex and VisiumHD 10x Genomics datasets

Cell Ranger processed output was first extracted to retain cells with mitochondrial read percentage of less than 25% and UMI counts greater than 500. Then we log-normalized the count matrix and performed PCA on the top 2,000 highly variable genes. We performed Leiden clustering to obtain the broad cell type categories. We then extracted the T cells and re-clustered the cells to identify granular cell populations.

We analyzed the VisiumHD data sets at 8 µm resolution. We first filtered and retained spots with mitochondrial reads percentage less than 25%, and UMI counts greater than 500. We then used Tacco^69^ to annotate the cell types in the Visium HD samples using the single-cell Flex data as reference. Since we have matched samples in both Flex and Visium, we used the identifical sample in Flex as reference, when performing the label transfer. We used the annotate() function in TACCO with parameters multi_center=3 and lamb=1e-3.

### Survival analysis

We ran BayesPrism^70^ to deconvolve the bulk RNA-seq data sets using the 10x Genomics single-cell flex as reference. This generated probabilistic predictions of deconvolution for each cell type in the reference atlas. We applied quantile cutoffs on the probabilistic predictions of IL-10^+^ and IL-10^−^ Tregs and generated Kaplan-Meier survival plots.

---

**Supplementary Figure 1.**
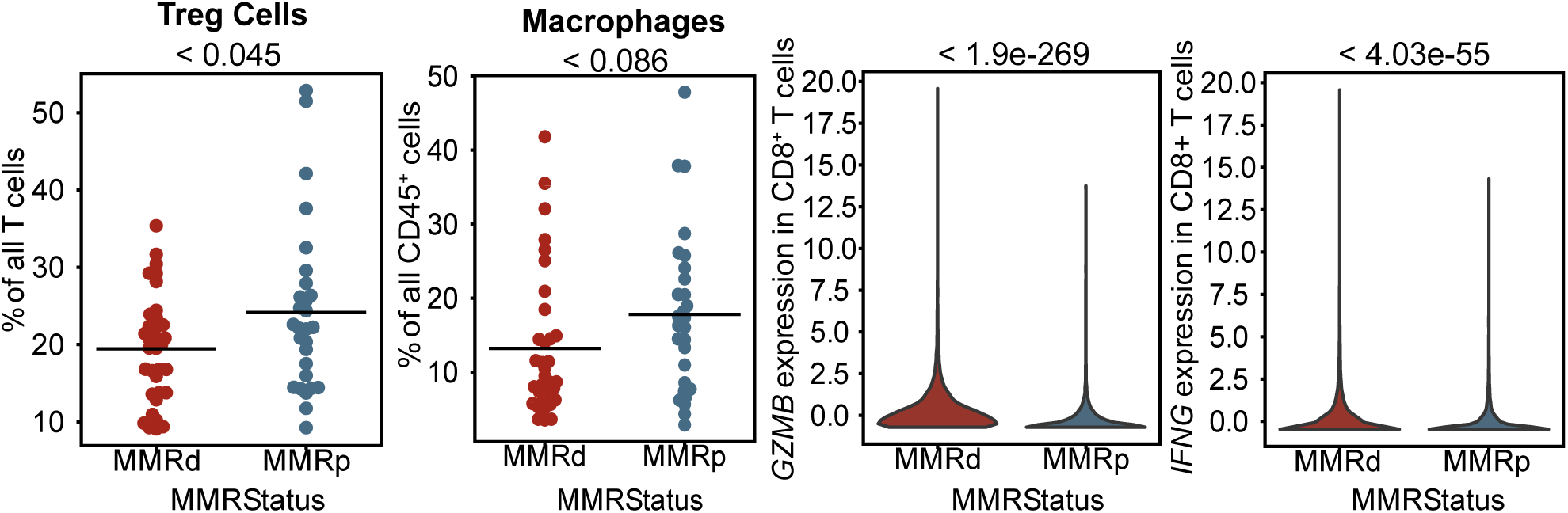
Distinct immune cell composition of human MMRd and MMRp CRC tumors. Reanalysis of scRNA-seq CRC datasets from Pelka et al.^24^ (**A**) Fraction of Tregs across patient samples grouped by MMRd and MMRp CRC. (**B**) Fraction of macrophage grouped by MMRd and MMRp CRC. (**C**) Differential *GZMB* and (**D**) *IFNG* expression in MMRd and MMRp CRC.

**Supplementary Figure 2.**
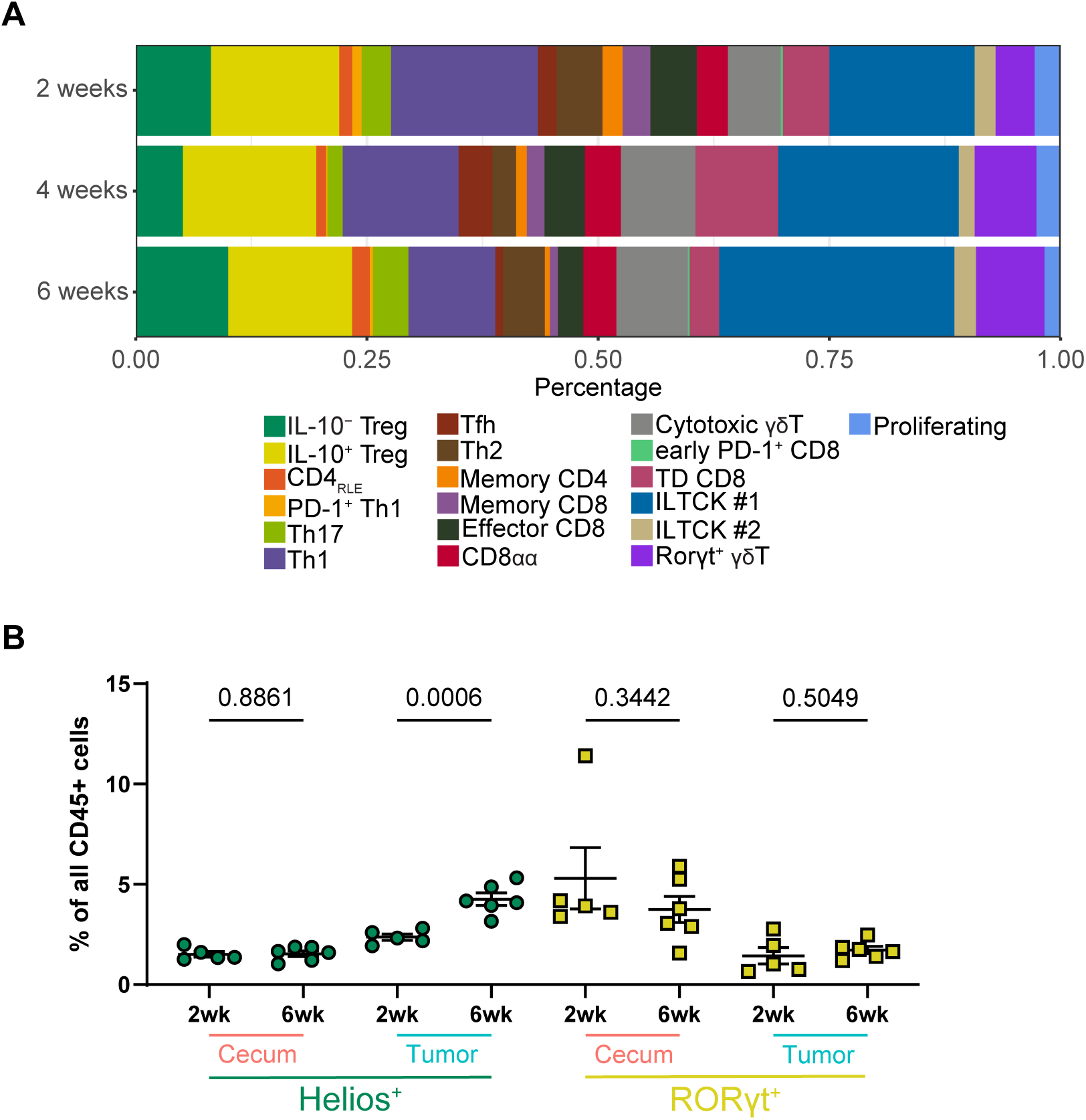
Temporal dynamics of heterogeneity of tumor and adjacent normal tissue T cells. (**A**) Analysis of T cell subsets in adjacent cecum at 2, 4, and 6 weeks after AKP organoid implantation based on scRNA+TCR-seq analysis (21,269 cells, 10,013 from normal adjacent cecal tissues and 11,256 from tumors). (**B**) Flow cytometric analysis of the frequencies of RORγt^+^ and Helios^+^ Treg cell subsets in tumor and adjacent normal tissue at the indicated time points after AKP tumor implantation. Pooled data from two independent experiments are shown.

**Supplementary Figure 3.**
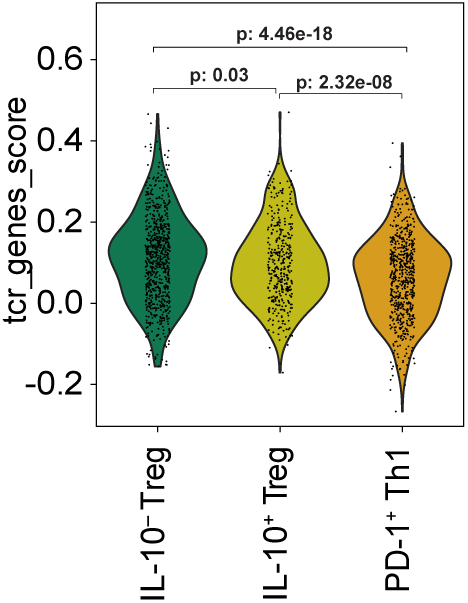
TCR- and IL-2-dependent gene expression in tumoral IL-10^+^ and IL-10^−^ Treg cell subsets. Violin plots of gene scores of genes associated with IL-2 signaling (n=30 genes) and TCR-signaling (n=30 genes).

**Supplementary Figure 4.**
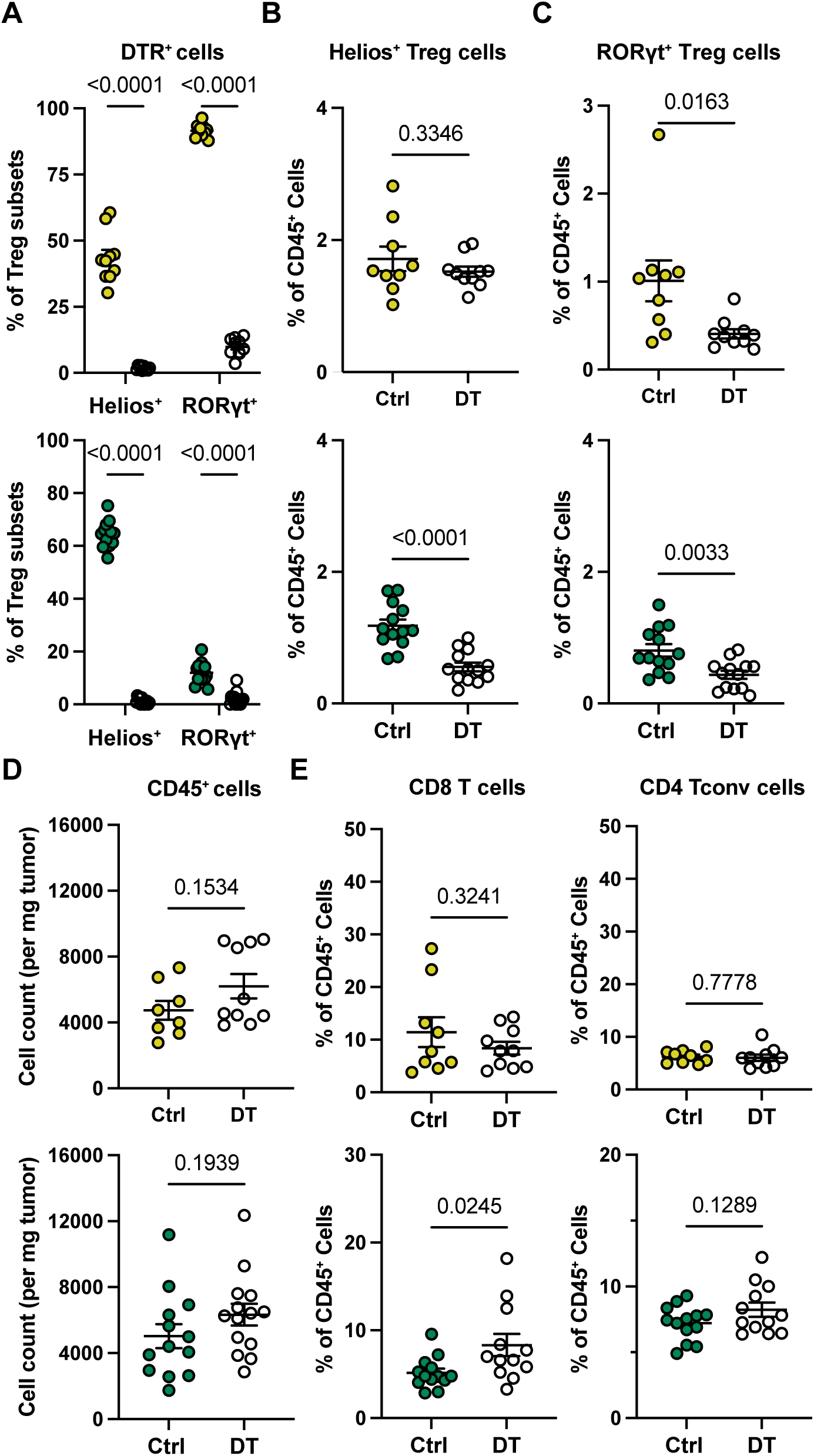
Selective ablation of IL-10^+^ and IL-10^−^ Treg cells does not alter total immune cell infiltration or CD8 and CD4 T cell composition of AKP tumors. (**A–E**) Flow cytometric analysis of the frequencies of indicated tumoral immune cell populations in DT and control (Ctrl) bDT-treated *Il10^tdTomato-Cre^ Foxp3^lsl-DTR^* (top) and *Il10^tdTomato-Cre^ Foxp3^flox-DTR^* mice (bottom). Pooled data from two independent experiments are shown.

**Supplementary Figure 5.**
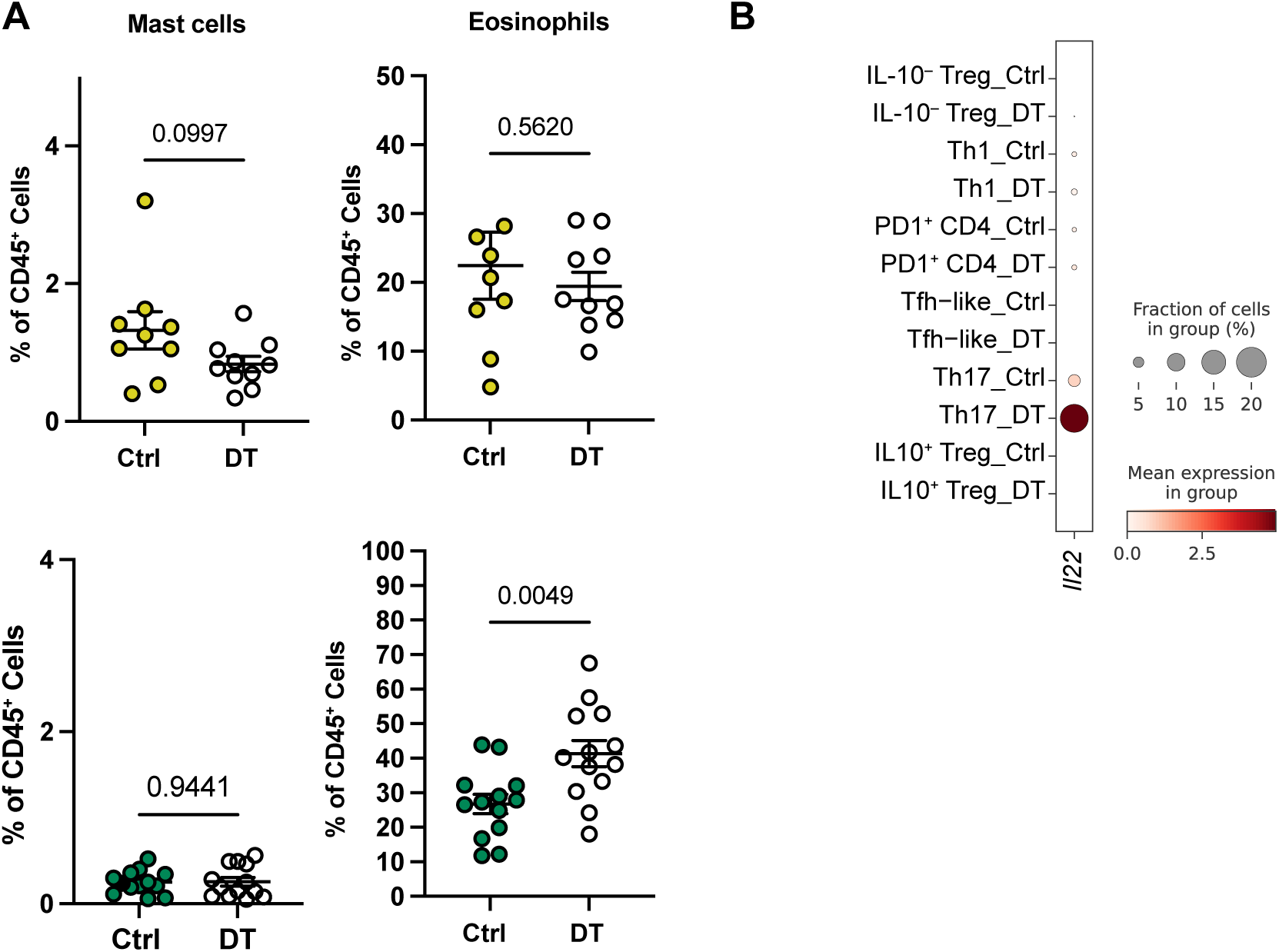
Selective IL-10^+^ and IL-10^−^ Treg cell loss results in distinct changes in myeloid cell and CD4 T cell composition of the AKP TME. (**A**) Flow cytometric analysis of mast cells and eosinophils in DT- and control (Ctrl) bDT-treated *Il10^tdTomato-Cre^ Foxp3^lsl-DTR^* (top) and *Il10^tdTomato-Cre^ Foxp3^flox-DTR^* mice (bottom). Pooled data from two independent experiments are shown. (**B**) *Il22* expression as measured by scRNA-seq analysis in indicated cell populations from mice.

**Supplementary Figure 6.**
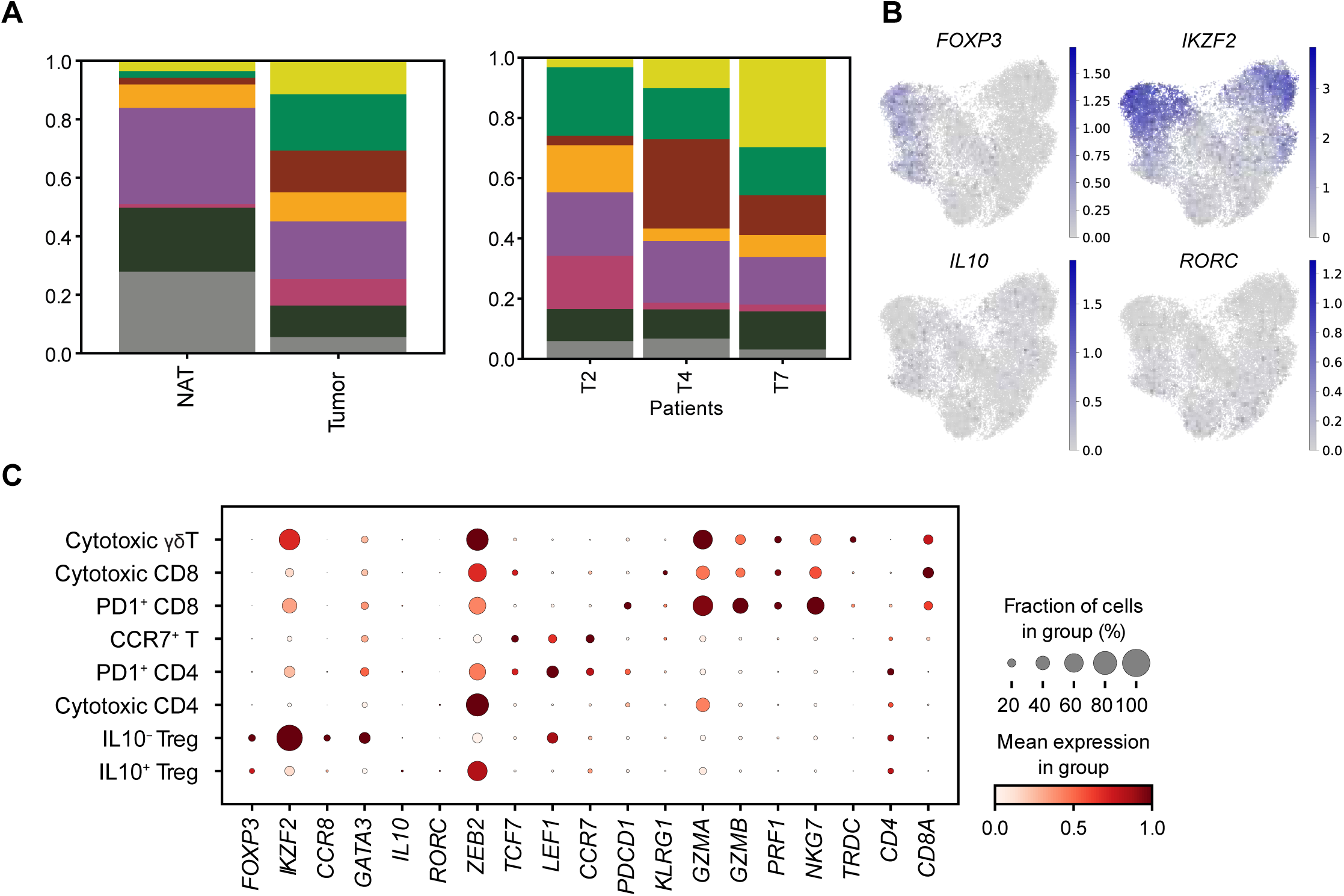
Single-cell multiome analysis of human CRC T cells. (**A**) Proportions of T cell subsets grouped by tissue (left) and by patients (right). (**B**) Expression of marker genes differentiating IL-10^+^ and IL-10^−^ Treg cells. (**C**) Dot plot of marker genes for T cell subsets.

**Supplementary Figure 7.**
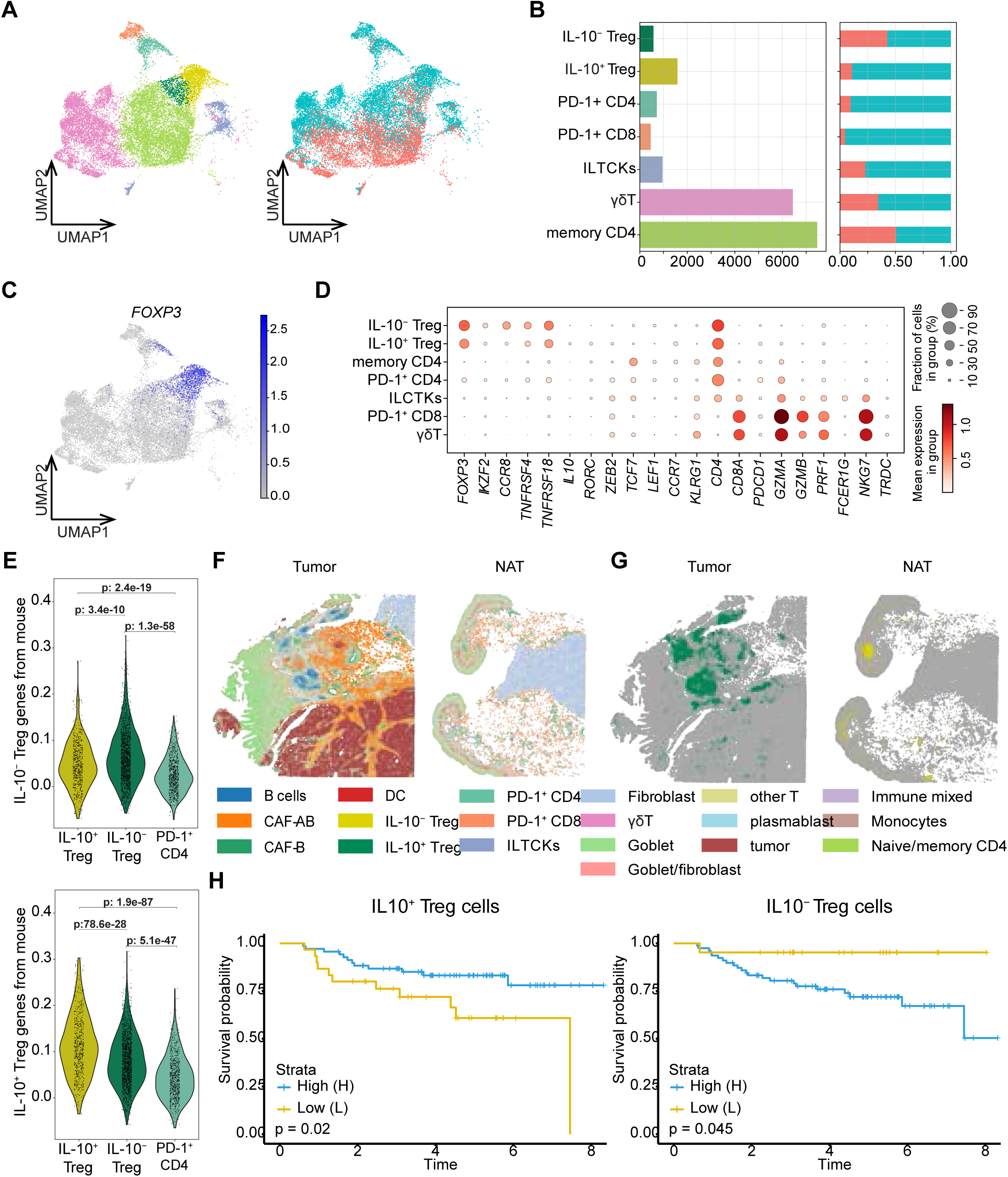
IL-10^+^ and IL-10^−^ Treg cell subsets in published single-cell RNA-seq and Visium HD spatial transcriptomic CRC datasets, and their association with survival probabilities. (**A**) UMAP of T cell subsets (n=18,198 cells) identified in 10x Genomics single-cell RNA-seq data. (**B**) Bar graphs depicting the number of cells for each cell type and the proportions for each tissue (cecum/tumor). (**C**) Expression of *FOXP3* to highlight the Treg cells. (**D**) Marker genes of T cell subsets. (**E**) Violin plots of gene scores calculated on the cells from single-cell flex using gene modules identified from mouse multiome for IL-10^+^ and IL-10^−^ Treg cells in Figure 3D. (**F**) Predictions of cell types in spatial 10x VisiumHD. The single-cell flex data was used as reference. (**G**) Spatial plots highlighting the predictions of IL-10^+^ and IL-10^−^ Treg cells in tumor and NAT. (**H**) Bulk RNA-seq from CRC patients (n=102) deconvolved into cell types from single-cell flex. Kaplan-Meier survival curves using quantile cutoffs on the prediction proportions of IL-10^−^ (left) and IL-10^+^ Treg cells (right).

**Supplementary Figure 8.**
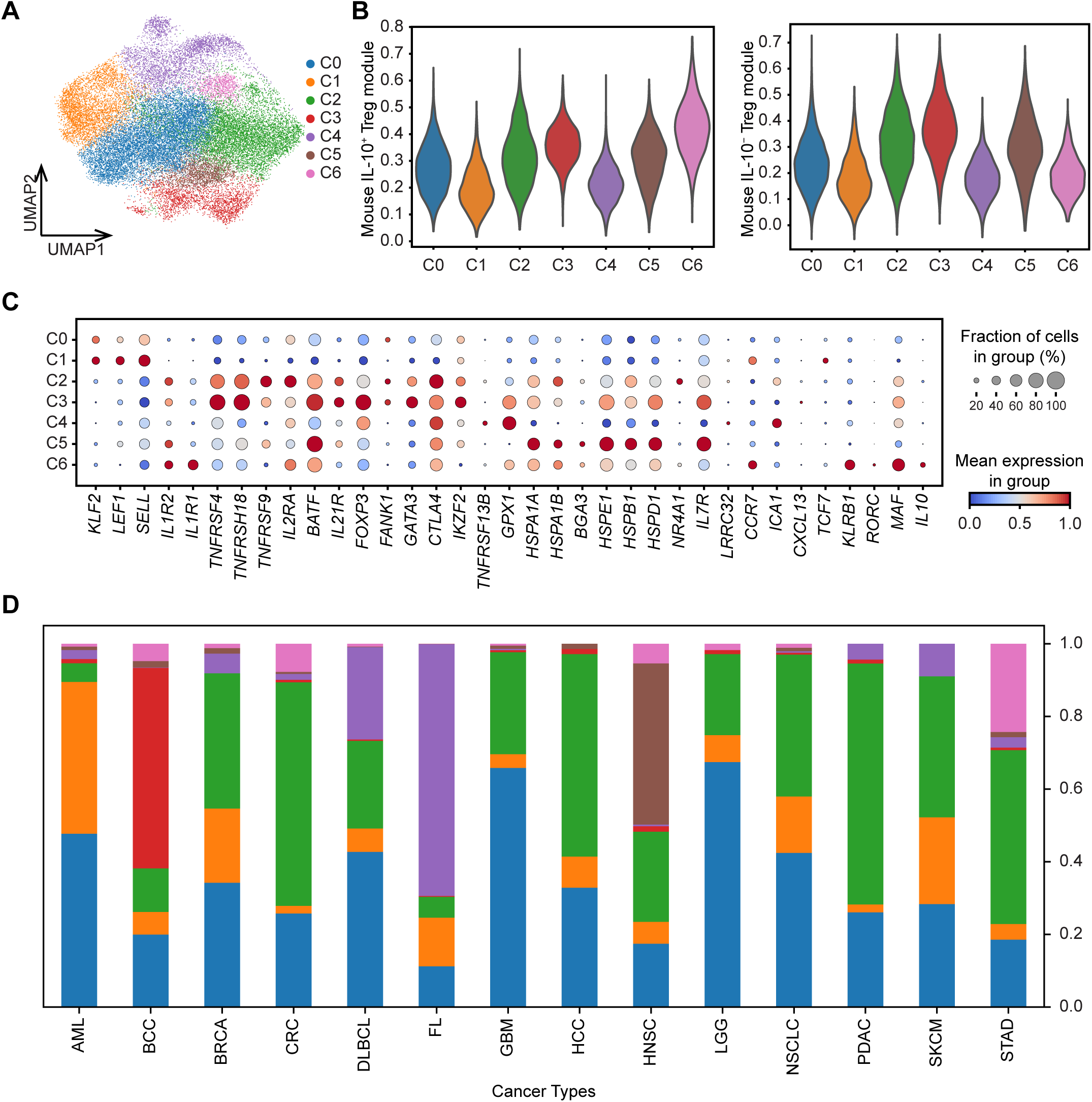
Pan-cancer analysis of human IL-10^+^ and IL-10^−^ Treg cell subsets. (**A**) UMAP visualization of previously identified Treg clusters from 14 tumor types, metastases, adjacent healthy tissue from cancer patients and PBMC, bone marrow, and lymph nodes from healthy donors (n=30,031 cells).^41^ (**B**) Gene scores of modules identified for IL-10^+^ and IL-10^−^ Tregs from CRC patient samples in Fig. 7F. (**C**) Markers of select genes differentiating the Treg clusters. (**D**) Composition of the Treg subsets across 14 cancer types.

